# Cellular Allostatic Load is linked to Increased Energy Expenditure and Accelerated Biological Aging

**DOI:** 10.1101/2022.02.22.481548

**Authors:** Natalia Bobba-Alves, Gabriel Sturm, Jue Lin, Sarah A Ware, Kalpita R. Karan, Anna S. Monzel, Celine Bris, Vincent Procaccio, Guy Lenaers, Albert Higgins-Chen, Morgan Levine, Steve Horvath, Balaji S Santhanam, Brett A Kaufman, Michio Hirano, Elissa Epel, Martin Picard

## Abstract

Stress triggers anticipatory physiological responses that promote survival, a phenomenon termed allostasis. However, the chronic activation of energy-dependent allostatic responses results in allostatic load, a dysregulated state that predicts functional decline, accelerates aging, and increases mortality in humans. The energetic cost and cellular basis for the damaging effects of allostatic load have not been defined. Here, by longitudinally profiling three unrelated primary human fibroblast lines across their lifespan, we find that chronic glucocorticoid exposure increases cellular energy expenditure by ∼60%, along with a metabolic shift from glycolysis to mitochondrial oxidative phosphorylation (OxPhos). This state of stress-induced hypermetabolism is linked to mtDNA instability, non-linearly affects age-related cytokines secretion, and accelerates cellular aging based on DNA methylation clocks, telomere shortening rate, and reduced lifespan. Pharmacologically normalizing OxPhos activity while further increasing energy expenditure exacerbates the accelerated aging phenotype, pointing to total energy expenditure as a potential driver of aging dynamics. Together, our findings define bioenergetic and multi-omic recalibrations of stress adaptation, underscoring increased energy expenditure and accelerated cellular aging as interrelated features of cellular allostatic load.

## Introduction

In response to environmental stressors, living organisms mount evolutionarily conserved responses that aim to increase resilience and promote survival, a phenomenon termed *allostasis*(*1, 2*). However, the chronic activation of these responses produces *allostatic load*(*3, 4*). Allostatic load is a multisystem state that reflects the added load, or “cost”, imposed by the activation of biological and physiological responses triggered by real or perceived stressors. Applied to humans, the allostatic load model predicts that when chronically sustained, allostatic load results in the maladaptive state of *allostatic overload*, which disrupts normal physiological functions and longevity(*5*). For example, while regular exercise improves glucose regulation and increases overall fitness(*6*), excessive exercise without appropriate recovery becomes damaging - disrupting glucose homeostasis and insulin secretion(*7*). Large-scale epidemiological studies also show that allostatic load, quantified by the abnormal levels of stress hormones, metabolites, and cardiovascular risk parameters, predicts physical and cognitive decline, as well as earlier mortality(*8–10*). These findings underscore the long-term, health damaging effects arising from chronically activated of stress pathways. However, the manifestations of allostatic load and overload at the cellular level, and whether they can accelerate cellular aging in a cell-autonomous manner, have not been fully defined.

Allostasis involves anticipatory processes that prepare the organism for potential threats - but it comes at a cost. Every cellular allostatic process - gene expression and protein synthesis, enzymatic reactions, hormone biosynthesis and secretion, and many others - consume energy in the form of ATP(*11*). In human cells, ATP is produced with moderate efficiency by glycolysis (*J*_ATP-Glyc_) in the cytoplasm, and with highest efficiency by oxidative phosphorylation (OxPhos, *J*_ATP-OxPhos_) within mitochondria. But building the molecular machinery for glycolysis and OxPhos pathways also comes at a cost, which is substantially higher for OxPhos owing to the extensive proteome demands of mitochondrial biogenesis and maintenance (>1,000 proteins) relative to glycolysis (-10 proteins)(*12*). Thus, a precise balance of energy derived from both glycolytic and OxPhos pathways allows cells to maintain optimal energetic efficiency.

Beyond their role in energy transformation, mitochondria respond to stress mediators such as glucocorticoids(*13*) and also regulate multiple physiological processes that encompass allostasis(*14*). This dual role as energy producers that enable allostasis, and as targets of stress hormones, positions mitochondria as key mediators of cellular and whole-body stress responses(*15*). We previously outlined the functional and structural mitochondrial recalibrations, as well as the potential detrimental effects that may occur in response to chronic stress, known as mitochondrial allostatic load (MAL)(*16*). In brief, potential *cellular* recalibrations include changes in cell size, energy requirements sustained by differential reliance on energy production pathways, or in the number of mitochondrial DNA (mtDNA) copies required to sustain OxPhos activity, whereas *mitochondrial* recalibrations would include features such as the accumulation of mtDNA damage (mutations, deletions) and changes in gene expression for the OxPhos machinery(*16*). However, these hypothesized cellular recalibrations, in particular the total energetic cost of allostatic load and the potential downstream maladaptive consequences of allostatic overload, have not been defined in a human system.

A major evolutionary-conserved stress mediator in mammals is the hormone family of glucocorticoids (GC; cortisol in humans, corticosterone in rodents). GC signaling acts via the glucocorticoid receptor influencing the expression of both nuclear(*17*) and mitochondrial genomes(*18*). Physiologically, GC regulates energy metabolism by acting on the brain, liver, adipose tissue, and skeletal muscles, elevating circulating glucose and lipids to supply energetic substrates and ensure readiness for potential threats. In humans and animal studies, GC signaling is considered a major contributor to allostatic load and stress pathophysiology(*19*). Chronically elevated circulating GC is linked to brain atrophy(*20*), cognitive decline and increased risk of Alzheimer’s disease(*21, 22*), as well as increased risk of cardiometabolic diseases(*21*). Therefore, based on previous short-term *in vitro* GC stimulation studies(*13, 23*), and the fact that *any* allostatic process requires active, ATP-dependent molecular events, we hypothesized that chronic GC signaling would trigger energy-dependent recalibrations among multiple domains of cellular behavior and bioenergetics. Moreover, in line with evidence in human studies (*24–26*) we further hypothesized that prolonged stress exposure in cells would trigger maladaptive outcomes, including multiple cellular aging markers and reduced cellular lifespan.

To quantify the bioenergetic and cellular manifestations of allostatic load across the lifespan, we apply chronic GC stimulation to primary fibroblasts from three healthy donors and deploy a longitudinal, high-frequency, repeated-measures strategy(*27–29*). We define the cellular features of allostatic load in four main parts. In *Part 1*, we quantify the bioenergetic and mitochondrial recalibrations and associated gene expression changes (Figures 1-4). In *Part 2*, we examine secreted cell-free DNA and cytokine signatures that mirror findings in human stress and aging studies (Figure 5-6). In *Part 3*, we profile well-established aging markers including mtDNA instability, DNA methylation-based epigenetic clocks, telomere length, and the Hayflick limit (Figure 7-8). Finally, in *Part 4*, we experimentally manipulate mitochondrial OxPhos to gain insight into the potential causal link between bioenergetics, aging, and cell death (Figure 9). Together, our findings define how cells recalibrate in response to chronic GC stress, highlighting increased energy expenditure (i.e., *hypermetabolism*) and accelerated cellular aging as interrelated, cell-autonomous features of allostatic load.

**Figure 1.**
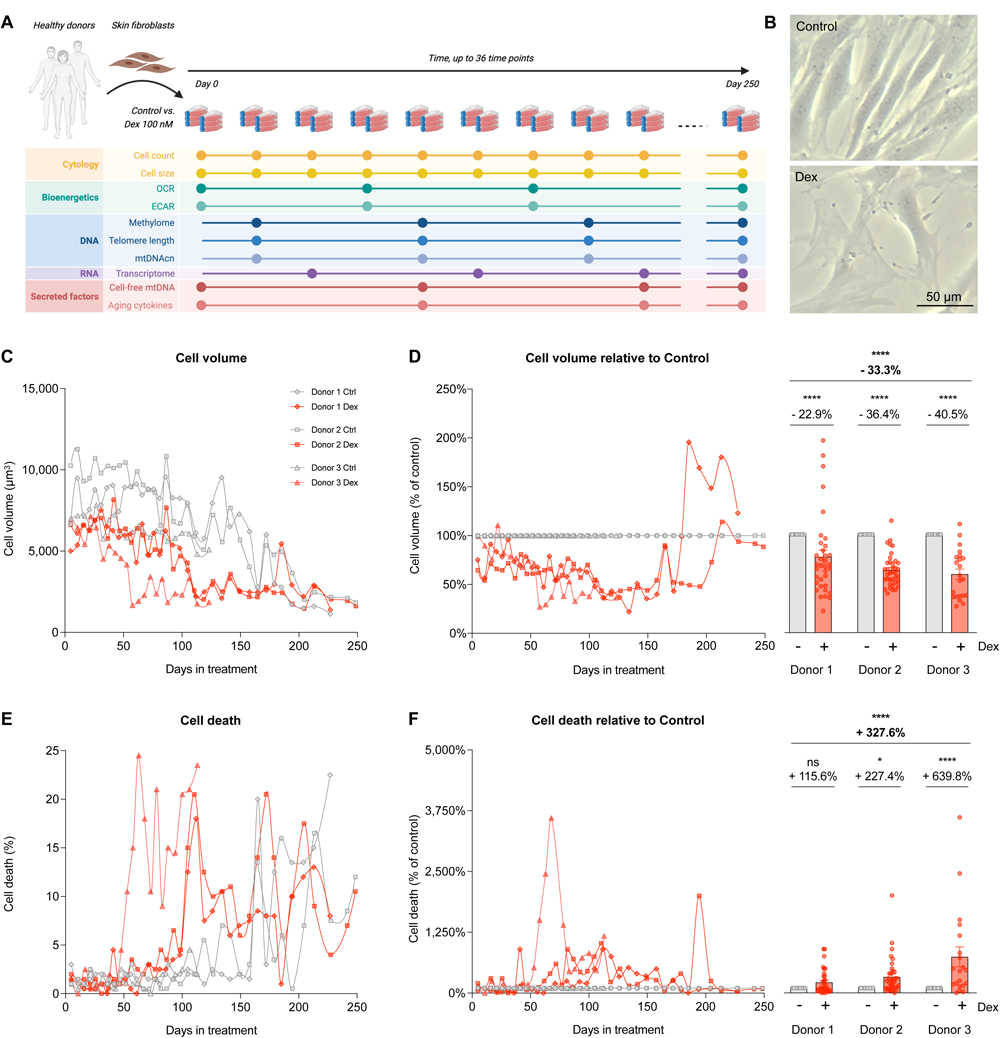
Longitudinal cytologic effects of chronic glucocorticoid signaling in primary human fibroblasts. (**A**) Study design: primary human fibroblasts derived from three healthy donors (Donors 1, 2, and 3) were cultured under standard conditions (control) or chronically treated with Dexamethasone (Dex, 100 nM) across their lifespan for up to 150-250 days, until replicative senescence. Cytologic parameters were evaluated every 5-7 days, while bioenergetics, DNA, RNA, and secreted factors parameters were evaluated every 10-20 days. (**B**) Representative images of untreated replicating control and Dex-treated cells from Donor 1 at 25 days of treatment. (**C**) Raw lifespan trajectories of cell volume. (**D**) To examine the effects of the chronic glucocorticoid signaling from the effects of aging, lifespan trajectories (left panel) and lifespan average effects (right panel) of Dex treatment on cell volume are expressed relative to the corresponding control time points for each donor. (**E-F**) Same as C-D but for the proportion of dead cells at each passage. n = 3 donors per group, n = 26-36 timepoints per donor. Lifespan average graphs are mean ± SEM, two-way ANOVA. * p < 0.05, **** p < 0.0001, ns: not significant.

**Figure 2.**
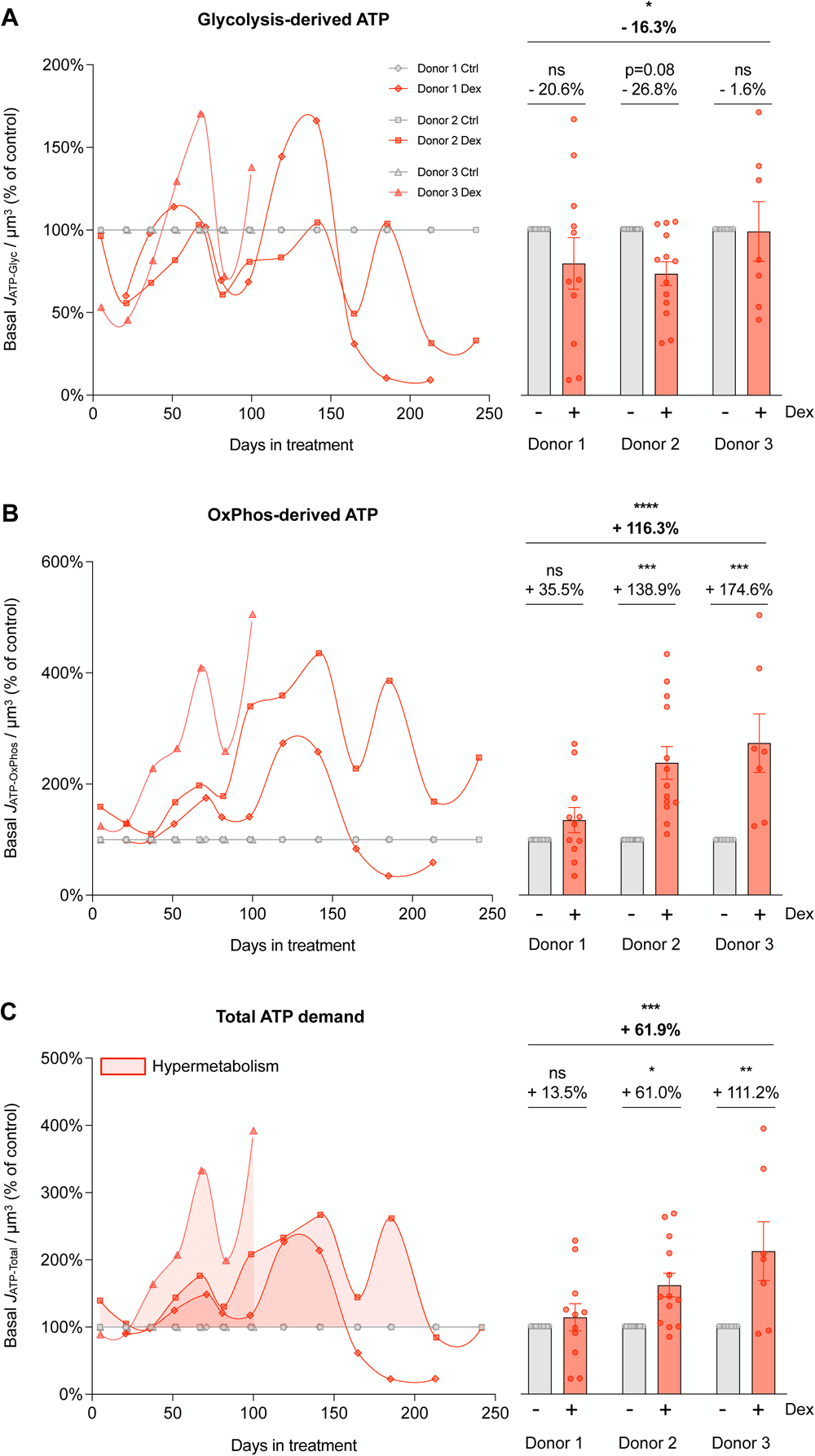
Cellular allostatic load is associated with hypermetabolism. **(A-C)** Energy expenditure trajectories across the cellular lifespan derived from Seahorse extracellular flux analyzer described in detail in ***Supplementary Material* Fig. S1**. (**A**) Lifespan trajectories (left panel) and lifespan average effects (right panel) of Dex treatment expressed relative to the corresponding control time points for each donor on basal glycolysis-derived ATP (*J*_ATP-Glyc_), (**B**) OxPhos-derived ATP (*J*_ATP-OxPhos_), and (**C**) total ATP production (*J*_ATP-Total_) corrected for cell volume. n = 3 donors per group, 8-13 timepoints per donor. Lifespan average graphs are mean ± SEM, two-way ANOVA * p < 0.05, ** p < 0.01, *** p < 0.001, **** p < 0.0001, ns: not significant. *J*_ATP-Total_ is the algebraic sum of *J*_ATP-Glyc_ and *J*_ATP-OxPhos_.

**Figure 3.**
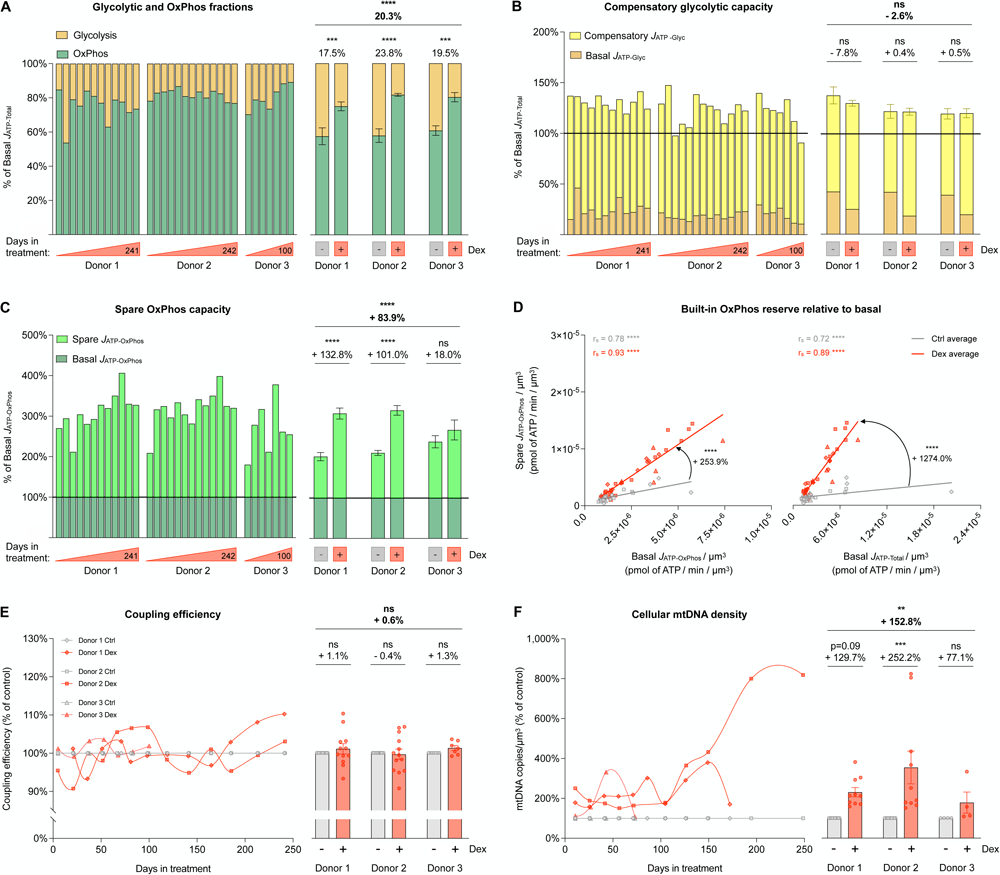
Cellular allostatic load involves a metabolic shift towards OxPhos. (**A**) Fraction of basal energy production from *J*_ATP-Glyc_ (yellow) and basal *J*_ATP-OxPhos_ (green) across the lifespan. (**B**) Compensatory glycolytic capacity, expressed as the percentage of basal *J*_ATP-Total_, achieved when OxPhos ATP production is inhibited with oligomycin. Basal *J*_ATP-Glyc_ levels are shown in dark yellow, and compensatory *J*_ATP-Glyc_ are shown in bright yellow. (**C**) Same as B but for spare OxPhos capacity, expressed as the percentage of basal *J*_ATP-OxPhos_ that can be achieved under uncoupled condition with FCCP. Basal *J*_ATP-OxPhos_ levels are shown in dark green, and spare *J*_ATP-OxPhos_ are shown in bright green. (**D**) Correlation between spare *J*_ATP-OxPhos_/cell volume and basal *J*_ATP-OxPhos_/cell volume (left panel) and between spare *J*_ATP-OxPhos_/cell volume and basal *J*_ATP-Total_/cell volume (right panel). (**E**) Lifespan trajectories (left panel) and lifespan average effects (right panel) of Dex treatment on coupling efficiency expressed relative to the corresponding control time points for each donor. (**F**) Same as C but for mtDNA copy number/cell volume. n = 3 donors per group, 8-13 timepoints per donor. Lifespan average graphs are mean ± SEM, two-way ANOVA. Correlation graphs show Spearman r and thick lines represent simple linear regression for each group. ** p < 0.01, *** p < 0.001, **** p < 0.0001, ns: not significant. *J*_ATP-Glyc_: ATP production rate derived from glycolysis. *J*_ATP-OxPhos_: ATP production rate derived from OxPhos. *J*_ATP-Total_: algebraic sum of *J*_ATP-Glyc_ and *J*_ATP-OxPhos_.

**Figure 4.**
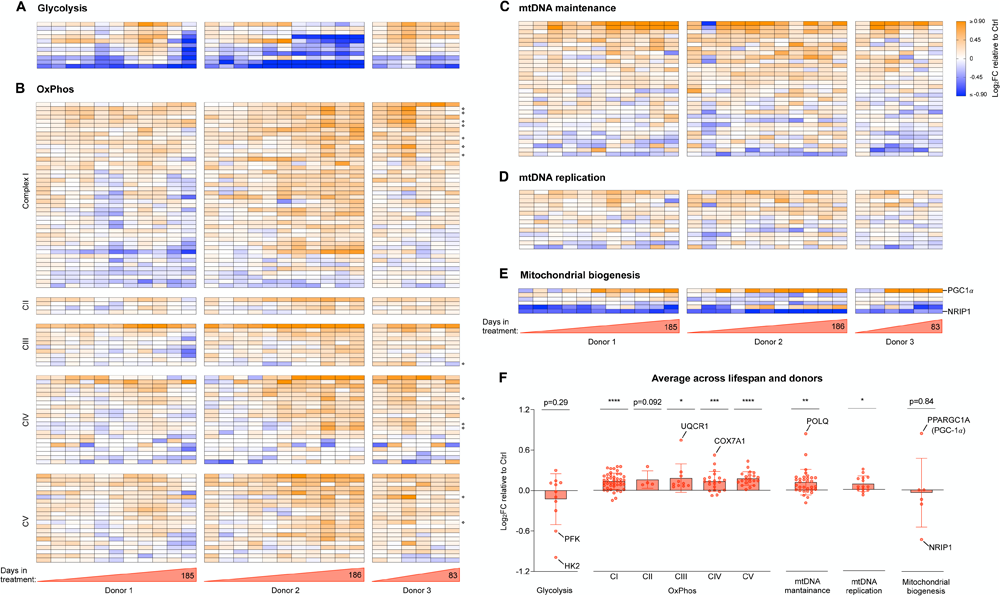
Cellular allostatic load involves transcriptional upregulation of OxPhos and mitochondrial biogenesis. (**A**) Heatmaps showing the effect of Dex treatment on the expression of glycolytic genes, expressed as the Log_2_ of fold change (Log_2_FC) of normalized gene expression relative to the corresponding control time points for each donor. (**B**) Same as A but for genes encoding the subunits of complexes I, II, III, IV and V of the OxPhos system, with mitochondrial-encoded genes marked with ❖ (**C-E**). Same as A and B but for genes encoding proteins associated with (**C**) mtDNA maintenance, (**D**) mtDNA replication, and (**E**) mitochondrial biogenesis. Two key regulators of mitochondrial biogenesis, one positive (PGC-1a) and one negative (NRIP1) are highlighted, showing expression signatures consistent with both activated and un-repressed mitochondrial biogenesis. (**F**) Average effect of Dex treatment shown in A-E. Each datapoint represents the gene average of the Log_2_FC values throughout the entire lifespan of the three donors (n=28 timepoints). Average graphs are mean ± SEM, with each gene shown as a single datapoint. One-sample t-test different than 0, * p < 0.05, ** p < 0.01, *** p < 0.001, **** p < 0.0001, ns: not significant. n = 3 donors per group, 9-10 timepoints per donor. Heatmap row annotation with individual gene names is provided in ***Supplementary Material* Fig. S4**.

**Figure 5.**
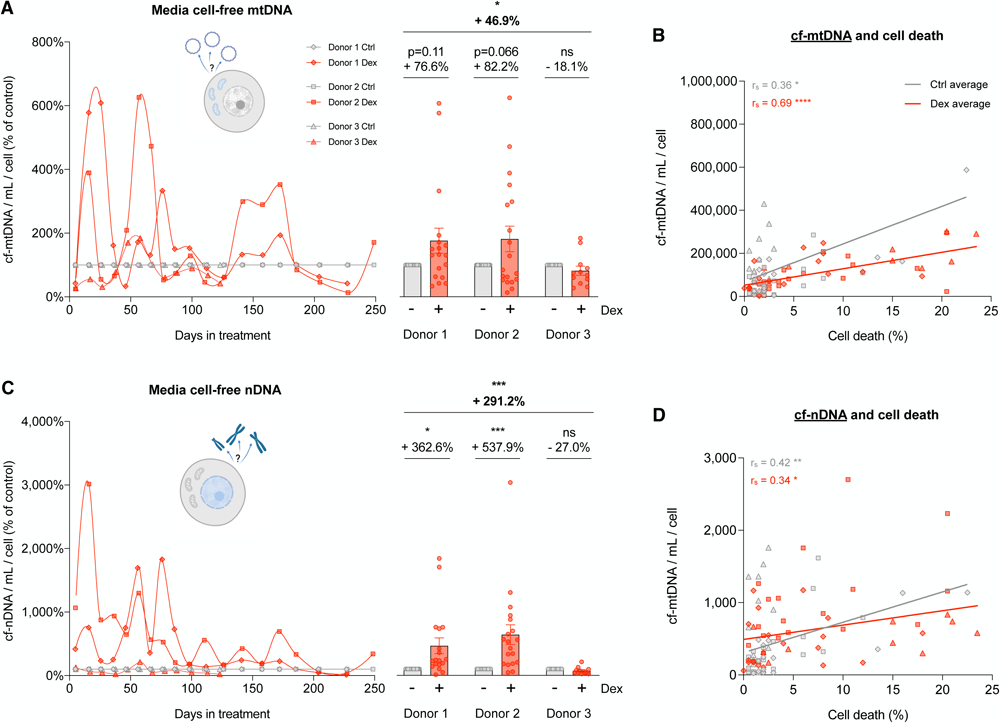
Cellular allostatic load increases cell-free DNA levels. (**A**) Lifespan trajectories (left panel) and lifespan average effects (right panel) of Dex treatment on cell-free mitochondrial DNA (cf-mtDNA) expressed relative to the corresponding control time points for each donor. (**B**) Correlation between cf-mtDNA and the proportion of dead cells at each passage. (**C**) Same as A but for cell-free nuclear DNA (cf-nDNA). (**D**) Same as B but for cf-nDNA. n = 3 donors per group, 6-10 timepoints per donor. Lifespan average graphs are mean ± SEM, two-way ANOVA. Correlation graphs show Spearman r and thick lines represent simple linear regression for each group. * p < 0.05, ** p < 0.01, *** p < 0.001, **** p < 0.0001, ns: not significant.

**Figure 6.**
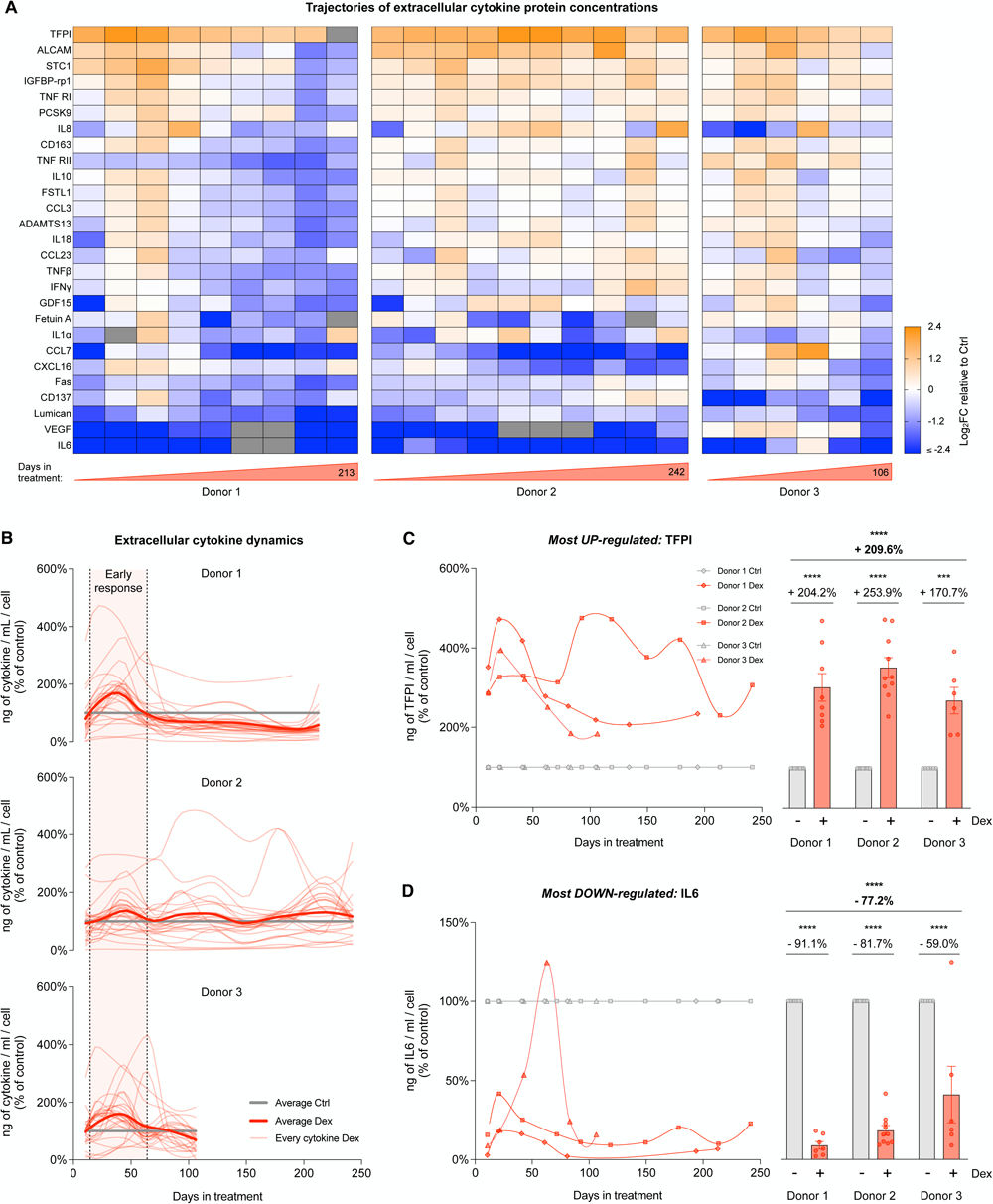
Cellular allostatic load alters cytokine release. (**A**) Heatmaps showing the effect of Dex treatment on the secretion of age-related cytokines, expressed as the Log2 fold change (Log2FC) of cytokine concentration (pg/mL of medium), relative to the corresponding control time points for each donor. (**B**) Lifespan trajectories of cytokine concentration in cells treated with Dex relative to the corresponding control time point for each donor. Thin curves in soft red represents individual cytokines; thick curves in red represent the average of all cytokines evaluated; thick lines in gray represent the control level. (**C**) Lifespan trajectories (left panel) and average effects (right panel) of Dex treatment on TFPII (most upregulated cytokine) levels per mL of culture media expressed relative to the corresponding control time point for each donor. (**D**) Same as C but for IL6 (most downregulated cytokine). n = 3 donors per group, 6-10 timepoints per donor. Lifespan average graphs are mean ± SEM, two-way ANOVA. *** p < 0.001, **** p < 0.0001, ns: not significant. Lifespan trajectories and average effects of Dex treatment on levels of every cytokine detected are shown in ***Supplementary Material* Fig. S7**.

**Figure 7.**
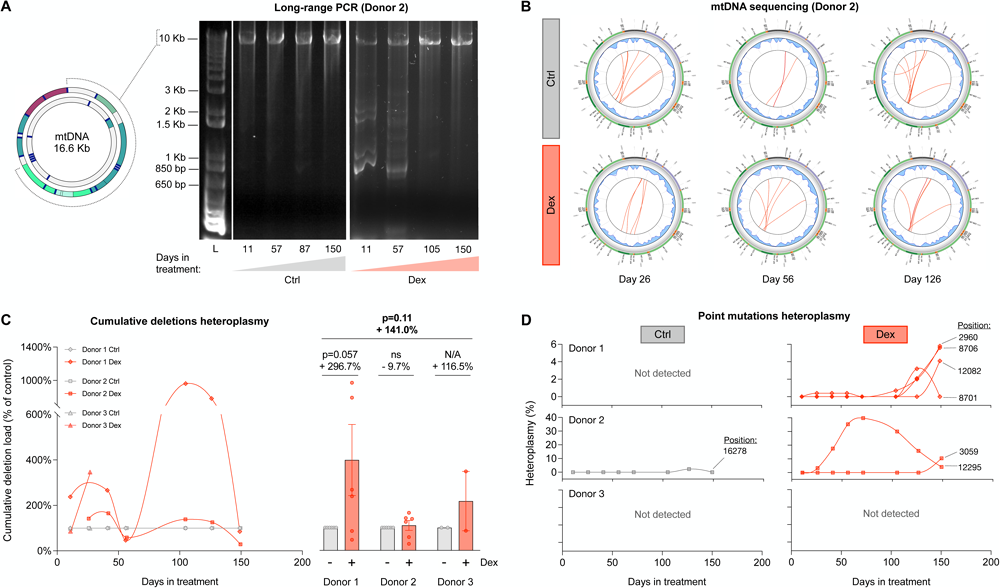
Cellular allostatic load causes mtDNA instability. (**A**) Long-range PCR (10 kb product) of mtDNA extracted from Donor 2 control and Dex-treated cells across lifespan, resolved by agarose gel electrophoresis. The presence of amplicons of sizes smaller than 10 kb reveal the presence of mtDNA molecules containing deletions. (**B**) Circos plots from mtDNA sequencing and eKLIPse analysis for mtDNA extracted from Donor 2 control and Dex-treated cells at days 26, 56 and 126. Each red line represents an individual deletion that spans the sequence contained between its two ends. The line thickness reflects the level of relative abundance or heteroplasmy for each deletion (see ***Supplementary Material* Fig. S8** for all donors). (**C**) Lifespan trajectories (*left* panel) and average effects (*right* panel) of Dex treatment on the cumulative heteroplasmy of all deletions present at a particular time point, expressed relative to the corresponding control time point for each donor. (**D**) Lifespan trajectories of individual point mutations heteroplasmy found in control (left panels) and Dex-treated cells (right panels) of the three donors. n = 3 donors per group, 2-8 timepoints per donor. Datapoints in lifespan trajectories are connected using the Akima spline curve, but the datapoints for Donor 3 in (C), for which due to insufficient time points the spline fit was not feasible and therefore datapoints were connected through a straight line. Lifespan average graphs are mean ± SEM, two-way ANOVA. ns: not significant, N/A: not applicable.

**Figure 8.**
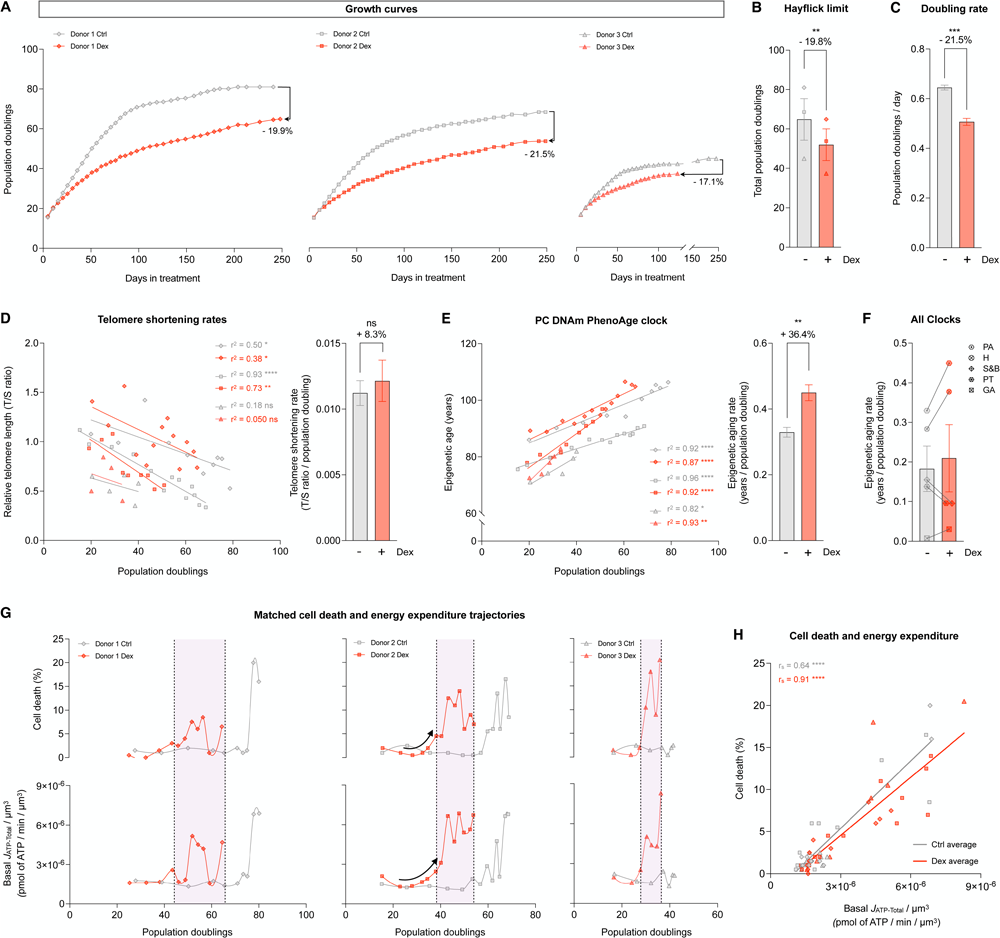
Cells under chronic allostatic load display accelerated cellular aging. (**A**) Lifespan trajectories of cumulative population doublings. (**B**) Hayflick limit for each donor of each group. (**C**) Group average early life doubling rate, inferred by linear mixed model of the population doubling trajectories within the first 50 days of treatment. (**D**) Telomere length across population doublings, with simple linear regressions for each donor of each group (left panel), and group average telomere shortening rate inferred by linear mixed model along the whole lifespan (right panel). (**E**) Epigenetic age calculated by the principal components (PC) PhenoAge epigenetic clock, with linear regressions for each donor of each group (left panel), and group average epigenetic aging rate inferred by linear mixed model along the whole lifespan (right panel). (**F**) Epigenetic aging rate for all the PC epigenetic clocks evaluated: PA: PhenoAge, H: Hannum, S&B: Skin and Blood, PT: PanTissue, GA: GrimAge. Detailed analysis of these epigenetic clocks is in ***Supplementary Material* Fig. S11** (**G**) Percentage of dead cells (upper panels) and basal *J*_ATP-Total_/cell volume (lower panels) across population doublings for Donor 1 (left panels), Donor 2 (middle panels) and Donor 3 (right panels). (**H**) Correlation between proportion of dead cells in every passage and Basal *J*_ATP-Total_/cell volume. n = 3 donors per group; timepoints per donor: n = 26-36 in A, n = 4-14 in D-E, n = 8-13 in G-H. Bar graphs are mean ± SEM, Satterthwaite method. Correlation graphs show Spearman r and thick lines represent simple linear regression for each group. * p < 0.05, ** p < 0.01, **** p < 0.0001, ns: not significant.

**Figure 9.**
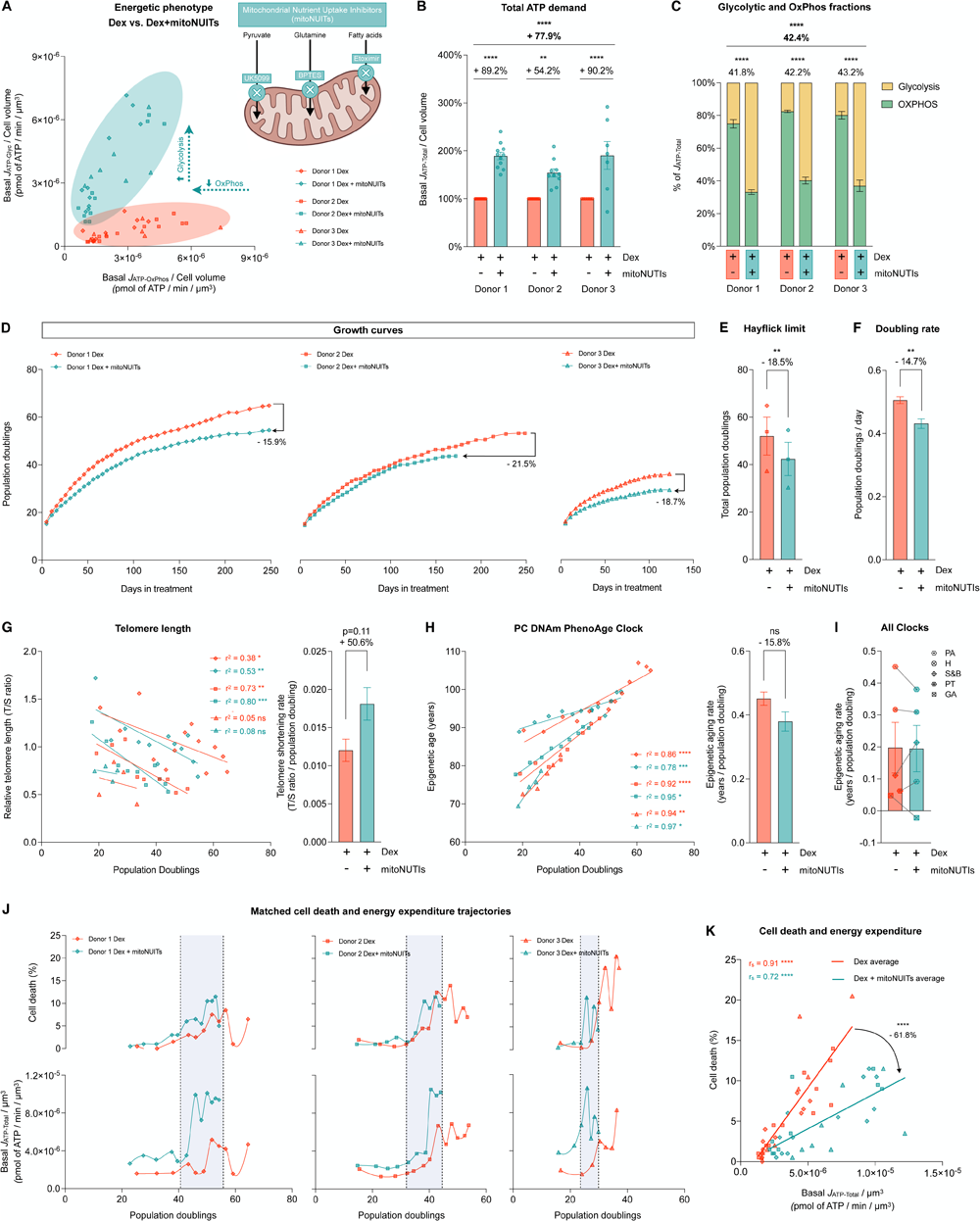
Hypermetabolism, not the metabolic shift towards OxPhos, predicts cell death. (**A**) Energetic phenotype of cells treated with Dex and Dex+mitoNUITs defined by Basal *J*_ATP-OxPhos_/cell volume and Basal *J*_ATP-Glyc_/cell volume across lifespan. (**B**) Lifespan average effects of Dex+mitoNUITs treatment on *J*_ATP-Total_/cell volume. (**C**) Values across lifespan (left panel) and average effects (right panel) of basal *J*_ATP-Glyc_ (yellow) and basal *J*_ATP-OxPhos_ (green) expressed as percentage of basal *J*_ATP-Total_. (**D**) Lifespan trajectories of cumulative population doublings. (**E**) Hayflick limit for each donor of each group. (**F**) Group averages of early life doubling rate, inferred by linear mixed model of the population doubling trajectories within the first 50 days of treatment. (**G**) Telomere length across population doublings, with simple linear regressions for each donor of each group (left panel), and group average telomere shortening rate inferred by linear mixed model along the whole lifespan (right panel). (**H**) Epigenetic age calculated by the principal components (PC)-adjusted PhenoAge epigenetic clock, with simple linear regressions for each donor of each group (left panel), and group average epigenetic aging rate inferred by linear mixed model along the whole lifespan (right panel). (**J**) Epigenetic aging rate for all the PC-adjusted epigenetic clocks evaluated: PA: Phene Age, H: Hannum, S&B: Skin and Blood, PT: Pan Tissue, GA: Grimm Age. (**K**) Percentage of dead cells (upper panels) and basal *J*_ATP-Total_/cell volume (lower panels) across population doublings for Donor 1 (left panels), Donor 2 (middle panels) and Donor 3 (right panels). (**H**) Correlation between proportion of dead cells in every passage and Basal *J*_ATP-Total_/cell volume. n = 3 donors per group; timepoints per donor: n=7-11 in B-C, n = 26-36 in D, n = 4-14 in G-H, n = 8-13 in J-K. Lifespan average graphs are mean ± SEM, two-way ANOVA in B, Satterthwaite method in E, F, H, and I. Correlation graphs show Spearman r and thick lines represent simple linear regression for each group. * p < 0.05, ** p < 0.01, *** p < 0.001, **** p < 0.0001, ns: not significant. mitoNUITs: Mitochondrial nutrient uptake inhibitors. *J*_ATP-Glyc:_ ATP production rate derived from glycolysis. *J*_ATP-OxPhos_: ATP production rate derived from OxPhos. *J*_ATP-Total_: algebraic sum of *J*_ATP-_ Glyc and *J*ATP-OxPhos.

## Results

### Chronic GC stress decreases cell volume and increases cell death

In *Part 1* of this study, we examine the cellular and energetic effects of chronic glucocorticoid stimulation. We treated human fibroblasts isolated from three healthy donors (Donors 1, 2, and 3, *see Methods* **Table 1**) with the GC receptor agonist dexamethasone (Dex, 100 nM) at a concentration known to trigger GC receptor translocation from the cytoplasm to the nucleus(*29*). To quantify allostatic load across the cellular lifespan (up to 150-250 days, depending on the lifespan of each cell line), cytologic parameters were evaluated every 5-7 days, in parallel with cellular bioenergetics, secreted factors, DNA-and RNA-based measurements every 10-20 days until cells reached replicative senescence, marked by a near absence of cell division (**Fig. 1A**). This design presents the major advantage of reducing potential bias from single-time points and to resolve potential time-dependent effects of chronic GC stress. Moreover, compared to similar experiments performed in immortalized cell lines, primary human fibroblasts derived from unrelated female and male donors also provide a more stringent test of generalizability for our conclusions. Throughout, data from Dex-treated cells of each donor is presented relative to the respective untreated control condition from the same donor.

**Table 1.** Primary human fibroblast information.

|  | Distributor | Catalog # | Genetic | Sex | Ethnicity | Age | Biopsy | Passage | Cell Line ID<sup>1</sup> |
| --- | --- | --- | --- | --- | --- | --- | --- | --- | --- |
| **Donor 1** | Lifeline Cell Technology | FC-0024 Lot # 03099 | Normal | Female | Caucasian | 18 years | Dermal Breast | 1 | HC2 (hFB13) |
| **Donor 2** | Lifeline Cell Technology | FC-0024 Lot # 00967 | Normal | Male | Caucasian | 18 years | Dermal Breast | 1 | HC1 (hFB12) |
| **Donor 3** | Coriell Institute | AG01439 | Normal | Male | Black | Newborn | Foreskin | 4 | HC3 (hFB14) |

<sup>1</sup>Refers to the ID given in the Cellular Lifespan Study Shiny Application (See *Data availability statement*)

Qualitatively, chronic Dex altered cell morphology, marked by a flattened appearance with more membrane protrusions and fewer contacts with surrounding cells (**Fig. 1B**). Dex caused a 25-50% reduction in cell volume within the first 5 days of treatment, which persisted across most of the cellular lifespan in all three donors (**Fig. 1C**), resulting in an average 33% volume reduction across the study period (p<0.0001, **Fig. 1C-D**). Towards the end of the lifespan between 200-250 days, both Dex-treated cells and controls converged towards the same apparent minimal cell volume. Trypan blue-based measures of cell death showed no initial effect of Dex on cell death for the first -50 days (**Fig. 1E**). However, Dex subsequently caused a marked elevation in cell death, averaging 3.3-fold increase across all three donors (p<0.0001, **Fig. 1E-F**). Thus, GC stress in this primary human cell system triggered robust time-dependent allostatic effects, including a reduction in cell volume and survival.

### Chronic GC stress triggers hypermetabolism

Survival and growth can be limited by energy constraints, meaning that the finite amount of cellular energy available limits various aspects of normal cell behavior(*30–34*). Cellular energetic constraints could arise in two main ways: from an increase in energy expenditure (i.e., ATP consumption), or from a reduced capacity to produce ATP. To evaluate the effects of chronic Dex on total energy expenditure and ATP production capacity, we directly measured extracellular flux of oxygen and pH (Seahorse extracellular flux analysis) longitudinally, every 10-28 days (***Supplementary Material* Fig. S1A**), using a validated mathematical approach to convert oxygen consumption rate (OCR) and extracellular acidification rates (ECAR) to ATP production rates(*35*). Under steady state, the ATP production rate is equal to ATP consumption rate, such that ATP synthesis rates can be interpreted as cellular energy expenditure. Furthermore, to obtain energy consumption per unit of cell mass (similar to measuring whole-body energy consumption per kg of body weight in humans), we normalized bioenergetic measures to the closest available measurement in our model, cell volume.

Using these strict parameters, we generated lifespan trajectories of three main parameters: i) resting (or basal) ATP production rates derived from glycolysis (*J*_ATP-Glyc_) and from OxPhos (*J*_ATP-OxPhos_), which reflects how much energy cells *consume* to sustain life and divide; ii) the maximal *J*_ATP-Glyc_ detected after pharmacologically inhibiting OxPhos, which reflects the glycolytic capacity to fulfill total basal cellular energetic demands; and iii) the maximal *J*_ATP-OxPhos_, which reflects the built-in spare OxPhos capacity above what is required to sustain cellular life (***Supplementary Material* Fig. S1B-C**).

Around day 25, Dex caused an initial decrease in basal *J*_ATP-Glyc_, which subsequently oscillated but remained on average 16% lower than control across the lifespan (p<0.05, **Fig 2A**). In contrast, Dex substantially increased basal mitochondrially-derived *J*_ATP-OxPhos_, which remained markedly elevated, reaching up to 3 to 4-fold higher rates than control in mid-life (p<0.0001, **Fig 2B**). Absolute *J*_ATP-Glyc_ and *J*_ATP-OxPhos_ values showed similar longitudinal behaviors as *J*_ATP-Glyc_ and *J*_ATP-OxPhos_ normalized by cell volume, with lifespan averages significantly lower and higher than controls, respectively (ps<0.0001, ***Supplementary Material* Fig. S2A-B).**

To quantify the total energetic costs of allostatic load in this system, we then calculated the combined basal ATP consumption rates derived from glycolysis and OxPhos. This total ATP consumption showed that Dex increased total resting energy expenditure across the lifespan (*J*_ATP-Total_/cell volume) by a striking 61.9% (p<0.001, **Fig 2C**). Dex-treated cells consume more energy per unit of volume to sustain life, reflecting *hypermetabolism*. This chronic state of hypermetabolism suggested the existence of costly allostatic recalibrations that may also involve changes in the degree to which cells use glycolysis or OxPhos to satisfy their elevated energy demand, which we investigated next.

### Chronic GC stress causes a metabolic shift towards OxPhos

To examine a potential shift in bioenergetics, we expressed *J*_ATP-Glyc_ and *J*_ATP-OxPhos_ as percentages of *J*_ATP-Total_ and compared Dex to control. Dex induced a significant shift towards mitochondrial OxPhos as the major source of ATP. Across the lifespan, Dex-treated cells derived 75-80% of their ATP from OxPhos, compared to 55-60% in control cells (p<0.0001, **Fig 3A**). Accordingly, *J*_ATP-Total_ in chronically stressed cells was strongly correlated to *J*_ATP-OxPhos_ (r_s_=0.92, p<0.0001, ***Supplementary Material* Fig. S2D**) and its variations along the lifespan were mostly explained by *J*_ATP-OxPhos_ (r^2^=0.87, p<0.0001, simple linear regression), highlighting the strong reliance of total energy expenditure on mitochondrial OxPhos.

This result led us to examine whether chronic GC stress directly impaired or inhibited glycolysis. When OxPhos is acutely inhibited with the ATP synthase (i.e., complex V) inhibitor oligomycin, mammalian cells naturally increase glycolytic activity to compensate for the missing *J*_ATP-OxPhos_ and fulfill the total cellular energetic demands(*36*). Here we found that after the addition of oligomycin, Dex-treated cells easily matched the basal *J*_ATP-Total_ of their control counterparts by upregulating *J*_ATP-Glyc_, and in fact sustain slightly higher than basal ATP consumption rates (ns, **Fig. 3B**). Thus, the intact glycolytic capacity of Dex-treated cells demonstrated that the metabolic shift towards OxPhos does not arise because of impaired glycolysis, but by upregulating the OxPhos system.

Accordingly, Dex increased the spare OxPhos capacity over that of untreated control cells by an average of 83.9% (p<0.0001, **Fig. 3C**). This means that for a given amount of energy required to sustain basal cellular functions, Dex-treated fibroblasts maintained an even larger spare *J*_ATP-OxPhos_ capacity than control cells - an effect that persisted across the lifespan (p<0.0001, **Fig. 3D**) and consistent with the anticipatory process of allostasis and allostatic load (*1–4*).

Such increases in OxPhos capacity could be explained by either increased improved mitochondrial respiratory efficiency or mitochondrial content. We first evaluated mitochondrial coupling efficiency, defined as the portion of oxygen consumption used towards ATP synthesis divided by the basal oxygen consumption rate. Dex did not change coupling efficiency, which oscillated by no more than 10% (within measurement error) from the control across the lifespan (n.s., **Fig. 3E**). Mitochondrial respiration efficiency therefore cannot explain the observed increase in the total energy expenditure nor the elevated OxPhos spare capacity. To indirectly monitor mitochondrial content, we next quantified total cellular mitochondrial DNA copy number (mtDNAcn) by qPCR, which showed that Dex induced approximately a doubling in mtDNAcn over the first 100 days of lifespan, followed by more substantial elevation at the end of life, particularly in one of the donors. On average across the lifespan, Dex-treated cells contained 936 mtDNA copies per cell, 97.8% higher than the control group (p<0.05, ***Supplementary Material* Fig. S3**). When accounting for the observed reduction in cell volume, cellular mtDNA density was 152.8% higher than in control (p<0.01, **Fig. 3F**), consistent with the increased reliance on OxPhos, the higher spare respiratory capacity, and the overall hypermetabolism in cells experiencing allostatic load.

### Chronic GC stress upregulates OxPhos and mitochondrial biogenesis gene expression

To examine the transcriptional recalibrations associated with stress-induced cellular allostatic load and hypermetabolism, we performed RNA sequencing (RNAseq) at 9-10 timepoints across the lifespan of each donor, and systematically queried the major glycolytic enzymes as well as OxPhos and mtDNA-related genes (**Fig. 4**, see ***Supplementary Material* Fig. S4** for individual gene labels). Consistent with the stable bioenergetic shift from glycolysis to OxPhos, chronic Dex downregulated most key glycolytic genes across the lifespan (**Fig. 4A**). This included the first enzyme in the sequence from glucose to pyruvate, hexokinase (HK2, -44.4%, p<0.001), as well as the rate limiting enzyme phosphofructokinase (PFK, -28.5%, p<0.001). On the other hand, chronic Dex upregulated most individual core subunits of the five OxPhos complexes. The most highly upregulated genes were UQCRC1 (complex III, +69.7%, p<0.001) and COX7A1 (complex IV, +52.2%, p<0.01), indicating a coordinated upregulation of the genetic OxPhos program (**Fig. 4B**).

Moreover, consistent with the elevated mtDNAcn in Dex-treated cells, genes encoding mtDNA maintenance-and replication-related proteins were both upregulated **(Fig. 4C-D**). This included the DNA polymerase theta (POLQ), which plays a key role in repairing double strand breaks by the theta-mediated end-joining mechanism(*37*) (+91.9%, p<0.001). This result also pointed to potential mtDNA instability as a feature of MAL(*16*) (see below, section “*Chronic GC stress causes mtDNA instabilit*y”). Moreover, Dex upregulated the master regulator of biogenesis PGC1a (peroxisome proliferator-activated receptor gamma coactivator 1-alpha, PPARGC1A(*38*)) by 106% across the lifespan, and simultaneously downregulated the *inhibitor* of biogenesis NRIP1 (nuclear receptor interacting protein 1, also RIP140(*39, 40*)) by 61.5% (ps<0.001, **Fig. 4E**). Together, the transcriptomics data summarized in **Fig. 4F** reveal a sustained stress-induced upregulation of the genetic programs involved in building additional OxPhos capacity and mtDNA copy number, consistent with the bioenergetic recalibrations and the anticipatory processes of cellular allostatic load in Dex-treated cells.

### Chronic GC stress increases cell-free mtDNA levels

In *Part 2* of this study, we turn our attention to the secretory profile of Dex-treated cells. In humans, two situations associated with allostatic load - acute psychological stress(*29, 41*) and aging(*42*) - are associated with increased circulating cell-free mtDNA. cf-mtDNA reflects the extracellular release of whole mitochondria or of mitochondria-free mtDNA(*43, 44*) and is an emerging mitochondria-derived inter-cellular signaling pathway(*45*). Motivated by the changes in mitochondrial OxPhos and mtDNA density in our system, we longitudinally investigated extracellular cf-mtDNA levels.

On average, cf-mtDNA levels across the lifespan were 46.9% higher in Dex-treated cells than controls (p<0.05) (**Fig. 5A**). In two of the three donors, chronic Dex strongly increased cf-mtDNA levels in the media, which oscillated and eventually matched cf-mtDNA levels observed in control cells by the end of lifespan. Previous work showed that cf-mtDNA release can be a selective and regulated process not primarily driven by cell death (reviewed in (*45, 46*)). In our model, cf-mtDNA was significantly correlated with the proportion of cell death at each passage (Dex r_s_=0.69, P<0.001, **Fig. 5B**), suggesting that a portion, although not all of the extracellular cf-mtDNA in fibroblasts under allostatic load is linked to increased cell death.

Dex also robustly elevated cell-free nuclear DNA (cf-nDNA, measured in parallel with cf-mtDNA), and showed a similar behavior to cf-mtDNA, with a strong increase along most of the lifespan for the same two donors followed by a gradual normalization. Dex-treated cells exhibited a lifespan average cf-nDNA 2-fold higher than controls (p<0.001, **Fig. 5C**). Similar to cf-mtDNA, cell death was significantly correlated with cf-nDNA but accounted for only a minor fraction (10-14%) of the variance in extracellular nuclear genome (Dex r_s_=0.34, p<0.05, **Fig. 5D**), suggesting that active processes other than cell death likely contributed to cell-free DNA release.

Interestingly, whereas media cf-mtDNA and cf-nDNA were strongly correlated in control cells (r_s_=0.90, p<0.0001), in response to chronic Dex, a large number of cf-mtDNA molecules were released at certain timepoints (up to 6-fold) without a corresponding elevation in cf-nDNA. This is reflected in a markedly lower correlation between cf-mtDNA and cf-nDNA in Dex-treated cells (r_s_=0.36, p<0.05, ***Supplementary Material* Fig. S5**). This pattern makes it unlikely that both genomes are systematically co-released in this model. Overall, these data define the temporally sensitive release of extracellular mitochondrial and nuclear DNA as a feature of GC-induced cellular allostatic load. Because energy must be consumed to fuel DNA synthesis and its extracellular release, these results point to cf-DNA release as a potential contributor to hypermetabolism.

### Chronic GC stress alters cytokine release

Considering that allostatic load in humans is characterized by elevated levels of neuroendocrine mediators(*47, 48*) as well as chronic inflammation(*49*), we next sought to investigate the effects of chronic Dex on the profiles of secreted cytokines. Using a custom-designed Luminex array of plasma cytokines associated with human aging(*50*), we detected a panel of 27 cytokines in the culture media of the three donors across the lifespan (**Fig. 6A**). While chronic Dex caused variable responses among different cytokines, it temporarily increased the secretion of multiple cytokine over the first 60 days, consistent with the classic inverted-U shaped stress responses(*13*) (**Fig 6B**). The peak cytokine response occurred around ∼30 days after the onset of GC stress, with a magnitude approximately double (i.e., ∼100% increase) for most cytokines relative to control cells. As for cf-DNA, because energy must be consumed to fuel cytokine/protein synthesis and their extracellular release, these results point to protein secretion as a potential contributor to hypermetabolism.

Further analysis revealed that Dex significantly altered the lifespan average secretion of 12 out of the 27 age-related cytokines. The strongest response was a stable elevation (average +210%) of Tissue Factor Pathway Inhibitor (TFPI, p<0.001, **Fig. 6C**), a cytokine related to the complement and coagulation cascades(*51*). The most strongly downregulated cytokine was Interleukin 6 (IL6, -77%, p<0.001, **Fig. 6D**), a pro-inflammatory cytokine well-known to be repressed by GC signaling(*52*).

Interestingly, when we queried the same cytokines at the transcript level using RNA sequencing, we observed a global downregulation in RNA levels among the three donors (***Supplementary Material* Fig. S6A-B**). In addition, the effect sizes across the lifespan were 2-3 times larger at the RNA level than at the extracellular protein level (***Supplementary Material* Fig. S6C-D**). Comparison of the secreted protein and transcriptomic data demonstrated that only half of the cytokines showed congruent (i.e., within 10% variation) responses (***Supplementary Material* Fig. S7**), highlighting a disconnect between quantitative measures of gene expression and cytokine release, and emphasizing the value of secreted protein quantification as an end phenotype. Thus, chronic GC signaling triggered a time-sensitive secretory phenotype, possibly involving non-transcriptional and/or non-genomic effects in the release of age-related cytokines.

### Chronic GC stress causes mtDNA instability

Thus far, we have examined key bioenergetic features of cellular allostatic load and related extracellular signaling behavior. In *Part 3* of this study, we turn our attention to potential maladaptive consequences of sustained allostatic load, namely evidence of allostatic overload. *In vitro* and *in vivo* studies have shown that chronic metabolic stressors, primary energetic defects, and aging result in mtDNA instability, which manifests as the accumulation of mtDNA defects(*53–57*). In Dex-treated cells, long-range PCR (LR-PCR) provided initial evidence that chronic glucocorticoid signaling may trigger mtDNA instability, as illustrated in multiple subgenomic bands consistent with mtDNA deletions, detected at different time points across the lifespan in two (Donors 1 and 2) out of three donors (**Fig 7A** shows data for Donor 2). We confirmed the accumulation of mtDNA deletions in all donors through deep mtDNA sequencing and quantified their relative abundance (i.e. proportion of mutant and normal mtDNA genomes, or heteroplasmy) across the lifespan using the eKLIPse pipeline(*58*) (**Fig 7B, *Supplementary Material* Fig S8A**). Dex induced a relatively large increase in total mtDNA deletion levels between days 20-50 of treatment among all three donors, suggesting that the effects of glucocorticoid stress on mtDNA instability may develop relatively rapidly. On average, Dex tended to increase the total mtDNA deletion burden by an average of 141.1% (p=0.11, **Fig 7C**). The absolute heteroplasmy levels remained low (0.02-0.33%) and therefore unlikely to have material effects on OxPhos capacity, as our data showed. This trend in mtDNA deletion load was not driven by an increase in the number of unique deletions, but rather by higher deletion heteroplasmy levels (***Supplementary Material* Fig S8B**). Furthermore, mtDNA deletions in Dex-treated cells were i) moderately larger in size, and ii) tended to cover the D-loop (***Supplementary Material* Fig S8C-E**), a region that physically interacts with the mtDNA maintenance proteins whose gene expression were upregulated (*see* Fig 4F).

Another marker of mtDNA instability is the accumulation of single nucleotide variants (SNVs, also known as point mutations). Along the lifespan, except for one low SNVs heteroplasmy in Donor 2 detected on day 125, control cells largely did not accumulate novel SNVs. In comparison, during the same portion of lifespan we identified 3 and 4 novel SNVs in Donors 1 and 2 (7 novel SNVs in total), one of which clonally expanded to reach 39.8% heteroplasmy (**Figure 7D**).

Replicating cultured cells accumulate few novel mtDNA deletions and SNVs(*59*), likely as a result of purifying selection (fibroblasts with deleterious mtDNA defects die and are eliminated from the cell population). Nevertheless, the apparent elevation in both spontaneous deletions and point mutations are consistent with the accumulation of damage as a feature of allostatic load, and collectively point to mtDNA instability as a manifestation of cellular, and specifically mitochondrial allostatic load (MAL).

### Chronic GC stress accelerates cellular aging

Combined, the mtDNA instability, premature cell size reduction, elevated cell death, as well as the early induction of most human age-related cytokines suggested that cellular allostatic load could impact aging trajectories. We first examined the growth curves for each donor, which reveal the maximal number of population doublings (i.e., cell divisions) that can be accomplished by each donor before reaching replicative senescence. This feature, also known as the Hayflick limit(*60, 61*), is the most closely related outcome to human longevity in this simple cellular model and could reflect allostatic overload.

Consistent with the notion that chronic stress accelerates cellular aging in humans (reviewed in (*62, 63*)), Dex caused a premature halting of population doubling, resulting in an average 19.8% reduction in the Hayflick limit across the three donors (p<0.05, **Fig. 8A-B**). This effect was associated with a decrease in cell division rate within the first 50 days of treatment (-21.5%, p<0.001, **Fig. 8C, *Supplementary Material* Fig. S9**). In the context of this substantial reduction in the division rate, and particularly given the reduced cell size in Dex-treated cells (-33% volume on average), the hypermetabolic phenotype of allostatic load becomes particularly striking. Considering cellular doubling rate and cell size, chronically stressed fibroblasts expend 107.9% more energy than control cells over the course of *each cell division* (p<0.001, ***Supplementary Material* Fig. S9**).

To examine the potential basis for the reduced cellular lifespan, we next examined two orthogonal measures of replicative senescence and cellular aging: telomere length(*24*), and DNA methylation-based epigenetic clocks(*64, 65*). We first quantified relative telomere length by qPCR across the lifespan for each donor and computed the rate of telomere shortening per cell division. Using a mixed effects linear model to compare rates of change over time, Dex showed an 8.7% non-significant acceleration in the telomere shortening rate (**Fig. 8D**); directly comparing the linear slopes of each Dex-treated cell line to its corresponding control showed an average acceleration of 27.6% across the three donors (p<0.05, paired t-test). Telomere length was also estimated through a DNA methylation-based algorithm(*66*). This measure documented an effect of similar magnitude towards accelerated shortening rate per population doubling (+17.0%, p<0.05, ***Supplementary Material* Fig. S10A-B**). These data thus provided converging evidence that chronically stressed cells experience more rapid telomere attrition during each event of genome replication, consistent with increased genomic instability and accelerated aging.

At the gene expression level, accelerated telomere attrition was associated with alterations in the expression of genes encoding telomere capping proteins. In particular, the major component of the shelterin complex, TPP1(*67*) was significantly downregulated across the lifespan for the three donors (-26.7%, p<0.0001), indicating that the telomeres in Dex-treated cells could be more vulnerable to DNA damage (***Supplementary Material* Fig. S10C-D**). On the other hand, several genes encoding the core components of the telomerase holoenzyme(*67*) were upregulated, including TERC, which was the most highly upregulated telomerase gene (+68.6%, p<0.0001, ***Supplementary Material* Fig. S10C-D**). The upregulation of TERC in the context of accelerated telomere shortening in Dex-treated cells is in agreement with clinical associations of chronic life stress and upregulation of telomerase activity in leukocytes(*68, 69*), possibly reflecting a (failed) attempt to reactivate telomerase to preserve or elongate telomeres, turning chronic allostatic load to overload.

Chronological age can also be predicted with high accuracy through DNA methylation-based algorithms, known as epigenetic clocks(*70–74*). Here, we generated a longitudinal DNA methylation dataset using the EPIC array(*28*), and deployed an approach that increases DNAmAge accuracy using a principal component-adjustment of the classic epigenetic clocks(*75*). The difference between the initial epigenetic age of the donors and their chronological age at the time of the biopsy can be explained the variable time and/or cell divisions underwent by each cell line before the beginning of the experiments, and because cells cultured *in vitro* experience an accelerated epigenetic aging rate compared to *in vivo* conditions (*76, 77*). Regardless of the estimated initial epigenetic age, by applying the clocks across the cellular lifespan, we can longitudinally quantify the rate of epigenetic aging relative to population doublings. The PC-PhenoAge clock(*74*) showed a significant 36.4% increase in the epigenetic aging rate (p<0.01, **Fig. 8E)**. A similar increase was found with the PC-Hannum clock (+33.1%, p<0.05), whereas non-significant effects were obtained using other clocks (Horvath Skin and Blood, Horvath Pan Tissue, and GrimAge, ***Supplementary Material* Fig. S11)**. Results across these five different epigenetic clocks were heterogenous but indicated that chronic GC stress accelerated the rate of epigenetic aging in 3 out of 5 clocks (Fig. **8F**). Together, these telomere and DNA methylation data provided converging evidence in primary human cells experiencing allostatic overload that hypermetabolism is linked to the rate of cellular aging.

### Age-related hypermetabolism and cell death occur near-synchronously

Given the premature age-related rise in cell death (*see* Fig. 1) and the profound mitochondrial and bioenergetic recalibrations associated with increased energy consumption in Dex-treated cells (*see* Fig. 2 and Fig. 3), we leveraged the longitudinal nature of our dataset to examine whether cell death was temporally related to hypermetabolism. Plotting both mortality (% cell death) and total energy expenditure (*J*_ATPtotal_/cell volume) relative to population doublings demonstrated that chronic Dex caused a left shift in the mortality curves, shifting the trajectory of each donor towards an earlier onset and progression along their lifespan (**Fig. 8G**, *upper* panels). Strikingly, Dex caused a similar and near synchronous left shift in energy expenditure trajectories, reflecting an early onset, hypermetabolic state that develops over weeks (**Fig. 8G**, *lower* panels). Comparing both mortality and energy expenditure at matched timepoints revealed a strong temporal correlation across the entire lifespan, among both control and Dex-treated cells (r_s_=0.64 and r_s_=91, respectively, ps<0.001, **Fig. 8H**), where 75-84% of the variance in cell death can be explained by the basal energy expenditure (i.e., how much energy cells must consume to sustain allostatic load), and vice-versa. In line with results linking the rate of telomere shortening and lifespan across species(*78*), these results underscore the close temporal association between the energetic cost of allostatic load (i.e., hypermetabolism) and possibly the ultimate consequence of allostatic overload, cell death or mortality.

### Hypermetabolism, not the metabolic shift towards OxPhos, predicts cell death

Previous work suggested that the shift from glycolysis towards OxPhos (e.g., during the differentiation of stem cells to “mortal” cell lineages) drives the susceptibility to detrimental age-related molecular alterations(*79*). Furthermore, mitochondria appear required for key cellular senescence features to manifest(*80*). Therefore, in *Part 4* of this study, we investigated whether in our model of cellular allostatic load the association of hypermetabolism with cell death and lifespan was specifically related to increased mitochondrial OxPhos activity (*J*_ATP-OxPhos_), or the total energy expenditure (*J*_ATP-_Total).

To disentangle these factors, we repeated the chronic Dex lifespan experiments while simultaneously downregulating OxPhos activity using a combination of inhibitors to block the mitochondrial import of the major carbon sources: pyruvate (UK5099(*81*)), fatty acids (Etomoxir(*82*)), and glutamine (BPTES(*83*)) (**Fig 9A**, ***Supplementary Material* Fig. S12A**). As expected, the mitochondrial nutrient uptake inhibitors (mitoNUITs) decreased OxPhos-derived ATP production in Dex-treated cells by an average of 21.2% (p<0.0001, ***Supplementary Material* Fig. S12B)**. This partial inhibition of OxPhos was associated with a supra-compensatory increase in glycolytic ATP production rates (+443.2%, p<0.0001, ***Supplementary Material* Fig. S12C**). Consequently, the Dex+mitoNUITs treatment induced an average increase in *J*_ATP-Total_ of 78.9% above the energy demand in Dex-treated cells (p<0.0001, **Fig. 9B**). Thus, compared to chronic Dex alone, Dex+mitoNUITs successfully suppressed OxPhos, but triggered an even more severe state of hypermetabolism (sustained predominantly by glycolysis). Therefore, we reasoned that if the observed allostatic overload phenotype -accelerated aging and premature cell death - was driven by the enhanced energy flux through OxPhos, the addition of mitoNUITs should at least partially rescue it. On the other hand, if cellular allostatic overload was directly driven by the total energy expenditure and hypermetabolism, then Dex+mitoNUITs would aggravate the pro-aging effects of allostatic overload.

Consistent with the latter alternative, the Dex+mitoNUITs combination either exacerbated or did not alter the pro-aging effects of Dex. Compared to Dex alone, Dex+mitoNUITs reduced the Hayflick limit by a further 18.5% (n=3 donors, p<0.05, **Fig. 9D-E)**, which was associated with a decrease in comparable cell division rate within the first 50 days of treatment (-14.7%, p<0.01, **Fig. 9F**). Dex+mitoNUITs tended to increase telomere attrition rate per population doubling (+50.6%, p=0.11, linear mixed model; +19.2%, p=0.45, paired t-test between slopes, **Fig. 9G**). However, the DNAm clocks-based rates of aging were not different between Dex and Dex+mitoNUITS (**Fig. 9H and 9I**), pointing either to potential mechanistic divergences or to a ceiling effect already achieved with Dex for epigenetic clocks.

In relation to cell death, the Dex+mitoNUITs treatment combination further left-shifted both the mortality and energy expenditure trajectories towards an earlier onset along the lifespan (**Fig 9J**), indicating a trend towards increased cell death compared to Dex alone (+28.6%, p=0.09). Again, the temporal correlation between energy expenditure and death across the lifespan remained strong and significant in Dex+mitoNUITs-treated cells (r_s_=0.72-0.91, p<0.001, **Fig. 9K**). Thus, diverting energy flux from OxPhos and elevating hypermetabolism with Dex+mitoNUITs treatment further exacerbated allostatic overload and age-related outcomes, implicating hypermetabolism, rather than flux through OxPhos, as the main driver of chronic stress-induced cellular allostatic overload.

In summary, our findings define the effects of chronic GC stress on cellular, bioenergetic and molecular recalibrations in primary human fibroblasts, highlighting increased energy expenditure (i.e., hypermetabolism) and accelerated cellular aging as interrelated, cell-autonomous features of allostatic load (**Fig. 10**).

**Figure 10.**
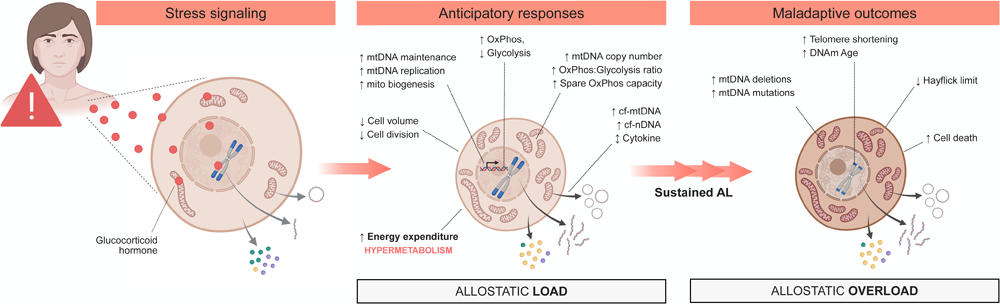
Summary diagram. Proposed model for the transduction of glucocorticoid signaling into cellular allostatic load and its interrelated cellular features, and the chronic downstream consequences of allostatic overload on cellular aging.

## Discussion

Although in humans the long-term effects of chronic stress on clinical outcomes such as cognitive and functional decline are well documented(*8, 9*), our understanding of how allostatic load manifests at the cellular level has remained incomplete. Our *in vitro* findings highlight hypermetabolism as a core manifestation of allostatic load, providing to our knowledge the first quantitative estimate of the energetic cost of allostatic load, increasing energy expenditure by >60% in this cellular model. Furthermore, our longitudinal data using well-established biological age indicators validated in humans, such as mtDNA instability, telomere length, and epigenetic clocks, also link hypermetabolism to accelerated aging biology as features of cellular allostatic overload. Lastly, stress induced-hypermetabolism was tightly linked to time of cell death, similar to the associations with mortality observed in humans. Thus, our results demonstrate that glucocorticoid signaling triggers energy-dependent allostatic recalibrations, which translates into age-related signs of allostatic overload. These results align with recent evidence of chronic stress pathophysiology in humans(*25*), and fill an important knowledge gap by defining the cellular and sub-cellular features of the original allostatic load model of chronic stress from McEwen and Stellar(*3*).

In healthy individuals, acute psychological stress, which is typically associated with elevated cortisol, increases whole-body energy expenditure by 9-67% (reviewed in (*84*)). Previous *in vitro* work in cultured rat neurons indicated that GC signaling could influence mitochondrial membrane potential over 72 hours(*13*), providing a basis for our first hypothesis that chronic GC stress could have long-term cellular bioenergetic consequences. Our cellular lifespan data allowed us to quantify the long-term, persistent cost of the anticipatory allostatic load. Our main finding is that it costs fibroblasts chronically exposed to Dex more energy (+62%) to sustain life, and remarkably more (+108%) to undergo each cell division. Plotting total energy consumption together with cell death also revealed a striking shift - potentially reflecting the transition from allostatic load to allostatic overload, where hypermetabolism and cell death were closely associated over time. Further work will be required to establish whether there is a causal link between hypermetabolism and fibroblast mortality. We note that relative to Dex, although the partial re-routing of metabolic flux to glycolysis with mitoNUITs did not rescue and in fact exacerbated the premature mortality phenotype (+28.6% mortality compared to Dex alone), it reduced the slope of the association between hypermetabolism and cell death (61.8%, p<0.001, *see* Fig. 9K). This relatively substantial shift may point to adaptive processes whereby cells are able to tolerate higher levels of hypermetabolism when deriving a portion of their energetic needs through both major pathways (OxPhos and glycolysis)(*12*), rather than through a single pathway. Nevertheless, our findings highlight the primary importance of total energy expenditure rather than of either pathway as a potential driver of allostatic overload.

Because our *in vitro* system consists of isolated cells, the observed effects must be cell-autonomous - meaning that they occur independent of inter-organ crosstalk, of the brain, and of other stress mediators encountered *in vivo*. This supports the idea that allostatic load is not a unique phenomenon of brain-bearing, complex multi-organ organisms. Instead, our findings point to allostatic load and allostatic overload as conserved processes able to manifest at the single cell level. Thus, the evolution of allostasis and allostatic load likely predated the evolution of the brain(*85*).

These results also begin to address potential mechanisms for how chronic GC stress causes hypermetabolism. Our data directly rules out three major potential confounders, namely cell volume, division rate, and mitochondrial OxPhos uncoupling. Larger and faster-dividing cells would require greater ATP demand. However, we found that chronically stressed cells under allostatic load were significantly smaller and divided significantly more slowly (possibly as an attempt to curtail the rising energetic cost arising from the demands of allostatic load). Within mitochondria, increased proton leak across the inner mitochondrial membrane, which uncouples respiration from ATP synthesis, can also drive cellular energy expenditure(*86*). But our data show that Dex-treated cells exhibit equal coupling efficiency than controls, ruling out this process as a cause of hypermetabolism in this system. Thus, our results implicate other active cellular processes as potential contributors for the increased “load” or energetic cost of living.

This state of hypermetabolism is likely driven not by a single process or signaling pathway, but by a multitude of inter-related processes. The major energetic costs within cells arise from gene transcription and translation(*87, 88*), from the maintenance of the plasma membrane potential, and from additional costs arising from protein (i.e., cytokine) secretion (*89*). The biogenesis of organelles is also expected to consume large amounts of ATP(*90*). This is particularly true for mitochondria, which contain large proteomes whose synthesis entails substantial energy expenditure and require genome replication(*90*). Furthermore, the heteroplasmic mixture of mutant and normal mtDNA molecules caused by mtDNA instability is predicted to directly increase the energetic cost of organelle maintenance(*91*). This cost may be particularly substantial in cells that upregulate mitochondrial biogenesis to build and sustain spare OxPhos capacity, well in excess of basal needs, as in our fibroblasts under allostatic load. Adding to these maintenance costs, our results revealed elevated extracellular secretion of cytokines and cf-mtDNA. Thus, the increases in mtDNAcn, the modest levels of mtDNA heteroplasmy, and secretion-related costs observed across the lifespan may all contribute to the increased energetic cost of living in Dex-treated fibroblasts. Additional work is required to determine the specific sequence of events whereby a neuroendocrine signal such as GC signaling leads to hypermetabolism, and the degree to which more complex stressors, such as chronic psychosocial stress, trigger hypermetabolism in whole animals including in humans (reviewed in (*84*)).

Another open question concerns the link between hypermetabolism and the reduced cellular lifespan that reflects allostatic overload. Our data cannot definitively establish directionality: does hypermetabolism cause the premature aging phenotype, or do accelerated biological aging processes drive increased energy expenditure? Nevertheless, to begin addressing this question we can consider two relevant bodies of literature: allometric scaling of metabolic rates and lifespan, and prospective studies of energy expenditure and mortality in humans.

First, there are well-defined relationships among body size, energy expenditure, and aging between animal species. These relationships show that animals of smaller sizes (e.g., mice, shrews) have correspondingly higher metabolic rates, age faster, and predictably live shorter lives than larger mammals (e.g., elephants)(*92–94*). Body weight and lifespan between animal species also scale linearly with the rate of telomere shortening, meaning that smaller animals with higher metabolic rates (i.e., relative hypermetabolism) exhibit correspondingly faster telomere shortening rates(*78*). These inter-species regularities, to which there are notable exceptions(*95, 96*), are at least superficially analogous to our human fibroblast data. Dex-induced allostatic load shifted cells towards a smaller cell size, correspondingly accelerated their mass-specific metabolic rates, accelerated the rate of telomere shortening and epigenetic aging, thus predictably resulting in shortened lifespan.

The second relevant body of literature prospectively links hypermetabolism, measured as elevated basal metabolic rate (BMR), with health outcomes and mortality in humans. Among healthy individuals, independent of well-recognized risk factors (e.g., age, body mass index, smoking, white blood cell count, and diabetes), hypermetabolism is associated with poor health in older individuals(*97*). In two other cohorts, elevated BMR also predicted earlier mortality over the subsequent 20-25 years(*98, 99*). Similarly, in patients with various illnesses (hepatitis B, amyotrophic lateral sclerosis, type 2 diabetes, and cancers), hypermetabolism predicts worse prognosis and mortality(*100–103*). Thus, we propose a model where hypermetabolism reflects the magnitude of allostatic load - i.e., how much energy cells and organisms are expending to maintain stable physiology. For reasons that still lack a molecular basis, hypermetabolism subsequently drives the accumulation of molecular damage and sub-cellular manifestations of allostatic overload, which accelerate biological aging and drive early mortality. This is recapitulated in cells and in patients with primary OxPhos defects, where genetically-or pharmacologically-induced hypermetabolism is associated with accelerated aging and shortened lifespan(*27*). The damaging nature of hypermetabolism may relate to general constraints on total energy flux imposing limits on molecular operations (i.e., Gibbs free energy dissipation rate (*104*)), or to intracellular tradeoffs between competing energy-consuming processes(*84*), although more work is required to understand how such constraints operate in human cells. One potential mechanism linking hypermetabolism and accelerated aging may involve oxidative stress, which is associated with psychosocial stress in humans (*105–107*), is induced by stress paradigms in animal models (*108–112*), and may play a role as a one of the hallmarks of aging (*113*). Further studies are required to establish the directionality, modifiability, and mechanistic basis of this stress-hypermetabolism-aging cascade.

In relation to aging biology, our study makes two noteworthy observations. First, it confirms the usefulness of replicative cellular lifespan models(*114–116*) with high temporal resolution sampling as an experimental approach to quantify the chronic effects of allostasis on a number of cellular, bioenergetic, and molecular outcomes. Moreover, deploying aging biomarkers validated in human populations to this *in vitro* system directly contributes to the interpretability of our findings, and to their potential physiological significance. Second, our relative telomere length data revealed that the loss of telomeric repeats is not a fixed quantity per genome duplication event (i.e., cell division), but that these can be decoupled by chronic stress exposure. The accelerated erosion of telomeric repeats per cell division caused by Dex implicate an effect of GC-mediated allostatic load on genomic stability, or on other aspects of telomere maintenance. Another study of Dex-treated human fibroblasts (one fibroblast line, aged for up to 51 days) reported no effect of cortisol or Dex on the rate of telomere shortening *per day in culture* (*117*). Our study (three donors, aged for up to 250 days) agrees with this observation when telomere shortening is expressed per day in culture. However, taking into account the reduced cell division rate reveals a markedly accelerated rate of shortening per event of genome replication, which we regard as the most relevant variable to understand telomere maintenance. Thus, future studies monitoring cell divisions over sufficiently long time periods have the potential to uncover connections between neuroendocrine stress exposure, hypermetabolism, and cellular aging.

Finally, some limitations of our study should be noted. While the epigenetic clocks can act as accurate molecular biomarkers of epigenetic age in several cell types and models, they still face challenges that make them less accurate in some models(*65*). Here, we used the PC clocks approach, which, by utilizing principal components rather than individual CpGs, stabilizes test-retest reliability and improves the age and age-related phenotypes prediction of classic DNAm clocks (*75*). In relation to our cellular model, compared to convenient immortalized cell lines, or to an experimental design that would include a single arbitrarily chosen donor, we noted substantial inter-individual variation among several measures between our three donors; Donor 1 was least affected, whereas Donor 3 was most affected on several variables. Unlike Donors 1 and 2 cells, which were obtained from the breast of healthy 18-year-old individuals, Donor 3 cells were obtained from the foreskin of a newborn who died prematurely shortly after birth (see *Methods* for details). This could have affected some of the basal cellular phenotype and responses to Dex, including the shorter lifespan of Donor 3. This experimental variability likely reflects true inter-individual differences. Thus, although this multi-donor design increases experimental variability, it importantly guards against the overfitting of results to a single donor/cell line, and therefore likely yields more robust and generalizable findings. In relation to the stressor, whereas chronic psychosocial stressors in humans involves the action of multiple hormones and metabolic factors, Dex is admittedly a simplistic model of stress. Therefore, although this targeted glucocorticoid stressor provides a strong proof-of-concept of the cell-autonomous energetic and molecular consequences of allostatic load, additional studies with other (combinations of) stress mediators are warranted. Finally, our analyses of the RNA sequencing and DNA methylation omics datasets are only partial, and more complete analyses of these data beyond the scope of the present manuscript could yield new insights into the global, longitudinal recalibrations and mechanisms underlying glucocorticoid-driven cellular allostatic load and the resulting allostatic overload. To examine these and related questions, the multi-omics dataset of chronic Dex is made available as a community resource (as well as other exposures, see *Data Availability Statement*)(*28*).

Finally, it is uncertain how the data in replicating cells reported here relates to cellular allostatic load in non-dividing, post-mitotic tissues. Our results documenting hypermetabolic, hypersecretory, and accelerated aging phenotypes is derived from actively replicating cells, which is similar to lymphoid and myeloid immune cells in the human body, but unlike the post-mitotic cells that populate several adult human tissues including the brain. We note that animal and human studies have demonstrated increases in whole-body energy expenditure in response to both acute (*118, 119*) and chronic stress (*120, 121*), collectively suggesting that stress-induced allostasis (cellular and systemic) are linked to hypermetabolism at the organism level (where most cells are post-mitotic). Moreover, the reported associations between perceived stress levels and the hallmarks of aging, including telomere shortening, DNA damage, mitochondrial impairments, cellular senescence, and inflammatory markers, appear to occur in both mitotic cells such as bone marrow leukocytes and thymocytes, as well as in post-mitotic brain tissue from different cortical and subcortical areas, including pre-frontal complex, amygdala and hippocampus (reviewed in (*122*)). Comparing metabolic and aging indicators longitudinally in both replicative and non-replicative cells could provide additional insights into the generalizable manifestations and mechanisms of cellular allostatic load.

In summary, we have defined multiple cell-autonomous features of GC-induced allostatic load, and mapped the long-term consequences associated with cellular allostatic overload in primary human fibroblasts. *First*, our work quantifies the added energetic costs of chronic anticipatory responses at the cellular level, thereby defining hypermetabolism as a feature of allostatic load. *Second*, we describe a hypersecretory phenotype that may contribute to hypermetabolism and mirrors findings from clinical studies. *Third*, we document manifestations of mitochondrial and cellular allostatic load including elevated mtDNA copy number and mtDNA density per cell, mtDNA instability, accelerated telomere shortening and epigenetic aging per cell division. In particular, we report a robust and specific temporal association between hypermetabolism and premature cell death, aligning with prospective observations in the human literature where hypermetabolism increases mortality risk. *Finally*, our experimental modulation of OxPhos and total J_ATP_ suggest that total energy expenditure, rather than flux through mitochondrial OxPhos, may have a particularly influential effect on cellular aging. Elucidating the mechanisms linking stress exposure, hypermetabolism, and shortened cellular lifespan will require further experimental work, as well as subsequent extension to well-controlled human studies. Resolving the cellular and energetic basis of allostatic load and chronic stress biology should reveal bioenergetic principles that can be leveraged to increase human resilience and health across the lifespan.

## Methods

### Human fibroblasts

Primary human fibroblasts of 3 healthy donors were obtained from certified distributors and described in detail in (*28*). A portion of lifespan data for the control (untreated) group was reported in (*27*). The characteristics of the three cell lines are summarized in Table 1. The ’age’ column indicates the age of the donor at the time of the tissue biopsy. During the course of the experiments, it came to our attention that Donor 3 was likely unhealthy since he had died prematurely of unknown cause 4 days after birth, which could have contributed to the variability in glucocorticoid responses and aging trajectories observed between the donors.

### Tissue culture

Cells were cultured under standard conditions of atmospheric O_2_, 5% CO_2_ and 37°C in DMEM (Thermo Fisher Scientific #10567022) supplemented with 10% Fetal Bovine Serum (FBS, Thermo Fisher Scientific), 50 µg/ml uridine (Sigma-Aldrich), 1% MEM non-essential amino acids (Thermo Fisher Scientific), and 10 µM-1.7 µM palmitate-BSA conjugate (Sigma-Aldrich). Treatment with dexamethasone (Dex, Sigma-Aldrich #D4902, 100 nM) and Dex plus the Mitochondrial Nutrients Uptake InhibiTors cocktail (Dex+mitoNUITs) began after 15-days of culturing post-thaw to allow the cells to adjust to the *in vitro* environment and were dosed every passage. The mitoNUITs cocktail included i) UK5099 (Sigma-Aldrich #PZ0160, 2 µM), an inhibitor of the mitochondrial pyruvate carrier (MPC) that interferes with the pyruvate import into the mitochondrial matrix ii) Etomoxir (Sigma-Aldrich #E1905, 4 µM), an inhibitor of carnitine palmitoyltransferase-1 (CPT-1) that interferes with the fatty-acid-derived Acyl-CoA import into the mitochondrial matrix; and iii) BPTES (Sigma-Aldrich #SML0601, 3 µM), an inhibitor of glutaminase 1 (GLS1) that prevents the conversion of glutamine to glutamate into the mitochondrial matrix. The combined action of these three compounds ultimately abates the availability of the tricarboxylic acid cycle (TCA) to produce Acetyl-CoA.

Cells were passaged every 5 ± 1 days through standard procedure using Trypsin-EDTA 0.25% (Sigma-Aldrich #T4049). Cell counts and cell volume assessment were performed using 0.4% Trypan Blue Stain and the Countess II Automated Cell Counter (Thermo Fisher Scientific #AMQAF1000). Total cell counts were used to calculate the doubling rate at each passage, and to determine the number of cells needed to reach ∼90% cell confluency by the time of the following passage. Cells not used for seeding or bioenergetic measurements were harvested and stored at -80°C for molecular analyses. Individual cell lines were terminated after exhibiting less than one population doubling over a 30-day period. The Hayflick limit was calculated as the cumulative number of population doublings of a cell line after termination. Brightfield microscopy images were obtained right before passaging using an inverted phase-contrast microscope (10X and 20X magnification, Thermo Fisher Scientific). The chronic Dex exposure experiments were performed once in each cell line. The reproducibility of the lifespan growth trajectories for each donor, and the reliability of specific outcome measures were established as described in (*28*).

Mycoplasma testing was performed according to the manufacturer’s instructions (R&D Systems) on media samples at the end of lifespan for each treatment and cell line used. All tests were negative.

### Bioenergetic parameters

The Seahorse XF Cell Mito Stress Test was performed in a XFe96 Seahorse extracellular flux analyzer (Agilent). ATP production rates derived from glycolysis (*J*_ATP-Glyc_) and oxidative phosphorylation (OxPhos, *J*_ATP-OxPhos_) were calculated from simultaneous measurements of cellular oxygen consumption rate (OCR) and extracellular acidification rate (ECAR) on a monolayer of 20,000 cells, with 10-12 replicates per group. The experimental conditions were, in sequence, as follows: i) 1 µM oligomycin-A, ii) 4 µM FCCP, and iii) 1 µM rotenone and 1 µM antimycin-A (Sigma-Aldrich). Raw OCR and ECAR measurements were normalized by a Hoechst-based cell count for each well (Cytation1 Cell Imager, BioTek), which provided more robust (i.e., less well-to-well variation) and stable results than results normalized to protein content. As illustrated in ***Supplementary Material* Fig. S1**, Basal OCR, basal ATP-linked OCR, proton leak OCR, maximal OCR, maximal ATP-linked OCR, compensatory OCR, coupling efficiency, basal ECAR, maximal ECAR, and compensatory ECAR were then utilized to derive ATP production rates from glycolysis and OxPhos as described previously by Mookerjee et al(*35*). OCR and ECAR measurements in the absence of glucose (0mM) or with 2-deoxyglucose (2-DG) were performed to confirm the specificity of the ECAR signal in control primary human fibroblasts under these culture conditions(*27*).

### mtDNA copy number

Cellular DNA was extracted using DNeasy Blood and Tissue Kit (Qiagen #69504). Duplex qPCR reactions with Taqman chemistry were used to quantify mitochondrial (mtDNA, ND1) and nuclear (nDNA, B2M) amplicons, as described previously(*123*). Primers and probes utilized were the following: ND1-Fwd: [5’-GAGCGATGGTGAGAGCTAAGGT-3’], ND1-Rev: [5’-CCCTAAAACCCGCCACATCT-3’], ND1-Probe: [5’-/5HEX/CCATCACCC/ZEN/TCTACATCACCGCCC/2IABkGQ/-3’]; B2M-Fwd: [5’-TCTCTCTCCATTCTTCAGTAAGTCAACT-3’], B2M-Rev: [5’-CCAGCAGAGAATGGAAAGTCAA-3’, B2M-Probe: 5’-/56-FAM/ATGTGTCTG/ZEN/GGTTTCATCCATCCGACCA/3IABkFQ/-3’]. Each sample was evaluated in triplicates. Triplicates with average Ct >33 were discarded. Triplicates with C.V. > 10% were also discarded. mtDNA copy number (mtDNAcn) was calculated as 2^βCt^ x 2, where βCt = average nDNA Ct - average mtDNA Ct.

### mtDNA deletions and point mutations

Long-range PCR (LR-PCR) was performed with mtDNA extracted as described in the mtDNA copy number section. Isolated DNA was amplified by PCR using Hot Start TaKaRa LA Taq kit to yield a 10-Kb product (Takara Biotechnology, #RR042A). Primers utilized were the following: Fwd: [5’- AGATTTACAGTCCAATGCTTC-3’], Rev: [5’-AGATACTGCGACATAGGGTG-3’]. PCR products were separated on 1% agarose gel electrophoresis in 1X TBE buffer, stained with GelGreen (Biotium #41005), and imaged using a GelDoc Go Imager (Biorad).

mtDNA next-generation sequencing was used to identify and quantify deletions and point mutations. The entire mtDNA was obtained through the amplification of two overlapping fragments by LR-PCR, using Kapa long Range DNA polymerase according to the manufacturer’s recommendations (Kapa Biosystems). The primer pairs utilized were tested first on Rho zero cells, devoided of mtDNA, to remove nuclear-encoded mitochondrial pseudogene (NUMTS) amplification. Primers utilized were the following: PCR1: Fwd: [5’-AACCAAACCCCAAAGACACC-3’], Rev: [5’- GCCAATAATGACGTGAAGTCC-3’] PCR2: Fwd: [5’-TCCCACTCCTAAACACATCC-3’], Rev: [5’- TTTATGGGGTGATGTGAGCC-3’]. PCR products were separated on a 1% agarose gel electrophoresis and NGS Libraries were generated using an enzymatic DNA fragmentation approach with Ion Xpress Plus Fragment Library Kit (Thermo Fisher Scientific). Sequencing was performed using an Ion Torrent S5XL platform with an Ion 540 chip, and signal processing and base calling were performed through the pre-processing embedded in house pipeline as described elsewhere(*124*). Demultiplexed reads were mapped according to the mtDNA reference sequence (NC_012920.1), and further analysis was performed through the eKLIPse pipeline(*58*) (https://github.com/dooguypapua/eKLIPse).

### RNA sequencing and transcriptomic analysis

Cells were lysed and stored at -80°C in TRIzol (Thermo Fisher Scientific). RNA isolation of all samples was performed as a single batch using a RNeasy Kit (Qiagen). All samples had an RNA integrity number (RIN) score >8.0, a A260/A280 ratio between 1.8-2.2, and no detectable levels of DNA. cDNA library was prepared from 1,500 ng of RNA using QIAseq FastSelect -rRNA HMR Kit (Qiagen) and NEBNext® Ultra™ II RNA Library Prep Kit for Illumina (New England Biolabs). cDNA was sequenced using paired-end 150 bp chemistry on a HiSeq 4000 System (Illumina) and yielded an average sequencing depth of 40 million reads per sample. Sequenced reads were aligned using the pseudoalignment tool kallisto(*125*) (v0.44.0) and imported using the ’tximport’ package(*126*) (v1.18.0). Variance stabilizing transformation (VST) was performed using the ’DEseq2’ package(*127*) (v1.30.1). Heatmaps and time courses show transcript levels as the Log_2_ of fold change (Log_2_FC) of normalized expression relative to the corresponding control time point for each donor. Categorized genes were selected using MitoCarta 3.0(*128*) and related literature on mitochondrial biogenesis(*37–40*).

### Cell-free DNA

At each passage, culture media was collected, centrifuged at 500g for 5 min, and the supernatant stored at -80°C until analyses until samples were processed as a single batch. To avoid cellular contamination, thawed media was further centrifuged at 1000g for 5 min, and the supernatant transferred to 96-well plates. Total cell-free DNA (cf-DNA) was isolated from 75 µL of cell culture media using an automated, high-throughput methodology based on the MagMAX Cell-Free DNA Isolation Kit (Thermo Fisher Scientific) that has been previously described(*129*). Duplex qPCR reactions with Taqman chemistry were used to simultaneously quantify mitochondrial (cf-mtDNA, ND1) and nuclear (cf-nDNA, B2M) amplicons, using the same primers and probes as described in the *mtDNA copy number* section. Serial dilutions of pooled human placenta DNA were used as a standard curve. mtDNA and nDNA copy number (copies/µL) of the standard curve samples were measured using singleplex, chip-based digital PCR (dPCR) on a Quant-studio 3D system (Thermo Fisher Scientific) according to the manufacturer’s protocol. The obtained values were then used to calculate the copy number of the experimental samples from the standard curve.

### Cytokines

Cytokine levels were quantified on the same culture media samples used for cf-DNA measurements. Two multiplex fluorescence-based arrays (R&D) were custom-designed with selected cytokines and chemokines whose levels in human plasma had been reported to be correlated with chronological age(*50*). Media samples were run along with negative controls (fresh untreated media), positive controls (healthy fibroblast aged for >200 days), and a standard curve following manufacturer’s instructions (Luminex). Cytokine concentrations were then normalized to the number of cells counted at the time of media collection to generate estimates of cytokine production on a per-cell basis.

### Relative telomere length

Relative telomere length was evaluated on the same genomic material used for other DNA-based measurements. Measurements were performed by single-plex qPCR reactions with SYBR Green chemistry and expressed as the ratio of telomere to single-copy gene abundance (T/S ratio), as previously described(*130, 131*). Primers utilized for the telomere (T) PCR were: tel1b: [5’- CGGTTT(GTTTGG)_5_GTT-3’], and tel2b: [5’-GGCTTG(CCTTAC)_5_CCT-3’]. Primers utilized for the single-copy gene (human beta globin) PCR were: Fwd: [5’ GCTTCTGACACAACTGTGTTCACTAGC- 3’] and Rev: [5’-CACCAACTTCATCCACGTTCACC-3’]. Each sample was evaluated in triplicates. Triplicates were subjected to Dixon’s Q test for outlier removal, and averaged values of T and S were used to calculate the T/S ratios. T/S ratio for each sample was measured twice. When duplicates showed C.V. > 7%, the sample was run a third time and the two closest values were used.

### DNA methylation and DNAmAge

Global DNA methylation was evaluated on the same genomic material used for other DNA-based measurements. DNA samples were submitted to the UCLA Neuroscience Genomic Core (UNGC) for bisulfite conversion and hybridization using the Infinium Methylation EPIC BeadChip kit (Illumina). DNA methylation data was processed in R (v4.0.2), using the ’minfi’ package (*132*) (v1.36.0). Data was normalized using functional normalization (Fun Norm), and RCP and ComBat adjustments were applied using the ’sva’ package (*133*) (v3.12.0) to correct for probe-type and plate bias, respectively.

Original DNAmAge was calculated using the online calculator (https://dnamage.genetics.ucla.edu/new) with normalization, using the age of the donor as the input age. This outputted the Horvath1 (i.e., PanTissue clock), Horvarth2 (i.e., Skin&Blood clock), PhenoAge, Hannum, and GrimAge estimated DNAmAges. More recent versions of the epigenetic clocks based on shared extracted variances among CpGs from principal components were also computed, yielding the PC-based DNAmAges for each clock (*75*). Stable estimates of the rate of epigenetic aging were obtained from the linear regression for each cell line between 27 and 210 days of growth, yielding three values per treatment condition.

### Statistical analyses

All statistical analyses were performed using GraphPad Prism (v9.0) and R (v4.0.2) using RStudio (v1.3.1056). All analyses were restricted to matching timepoints between control and the Dex or Dex+mitoNUITs treatments from the Cellular Lifespan dataset(*28*). In graphs of lifespan trajectories, datapoints are connected using the Akima spline curve. Lifespan average graphs show each passage as a single datapoint, and bars represents mean ± standard error of the mean (SEM). The percent differences between groups (i.e., Control vs Dex; or Dex vs mitoNUITs) was computed separately for each cell line (i.e., Donors 1, 2, 3), and groups were compared by two-way ANOVA treating each timepoint as a unique observation. Analyses of gene expression (i.e., HK2, PFK, UQCRC1, COX7A1, POLQ, PPARGC1A, and NRIP1, TPP1, TERC) were similarly performed by two-way ANOVA across all donors, without adjustment for multiple comparisons. To analyze temporal associations between variables, we performed non-parametric Spearman correlation, and report the correlation coefficient r_s_. To obtain proportions of shared variance between variable pairs, the squared coefficient of correlation from simple linear regression was calculated. To estimate the average expression of transcriptional pathways, the average Log_2_ fold change values of each gene were averaged across all timepoints of the lifespan of three donors, and p values obtained using a one-sample t-test against the reference value = 0 (i.e., no change in expression relative to control). For analyses of the Hayflick limit, a single value was obtained per donor (total n=3), and group averages were compared using ratio paired t-test (two-tailed). Because of the small sample sizes for these outcomes the p values are less reliable than the estimated differences (%) across multiple aging biomarkers. For analysis of the early-life doubling rate (i.e., first 50 days in treatment), doubling rate was modeled with linear mixed models using the lme4 package v1.1-26(*134*) with fixed effects for population doublings, treatment condition (Dexamethasone vs. control), and their interaction. We used random intercepts at the cell line level:

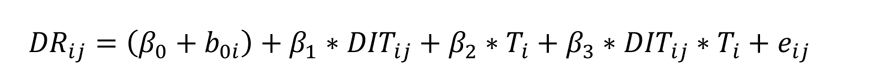

Where DR is doubling rate, i,j are cell line and measurement respectively, β and b denote fixed and random effects, DIT is days in treatment, T is treatment condition (0 = Control, 1 = Dexamethasone), and e is error. We assessed statistical significance using the Satterthwaite method implemented in the lmerTest package v3.1-3(*135*). Analyses of telomere shortening rate and epigenetic aging rate were performed using linear mixed models in the same way described above, but across population doublings instead of days in treatment. The significance threshold for all analyses was arbitrarily set at 0.05, and effect sizes are provided throughout.

## Supplementary Material

Supplemental Figures S1-S12 are available in the Supplemental Material.

## Author contributions

G.S. and M.P. designed experiments. G.S. performed cellular lifespan studies. A.S.M. performed quality control experiments. B.S.S. performed alignment and quality control of transcriptomic data. G.S. measured mtDNAcn. S.A.D. and B.K. measured cf-mtDNA. K.R.K. performed cytokine arrays and LR-PCR. C.B., V.P., G.L. performed mtDNA sequencing. J.L. and E.S.E. measured telomere length. S.H., A.H.C., and M.L. contributed the original and PC-DNAm clocks. G.S and N.B.A curated the database.

N.B.A. performed the data processing and prepared the figures. N.B.A. analyzed and interpreted the data with G.S. and M.P. N.B.A. drafted the manuscript with G.S. and M.P. All authors reviewed the final version of the manuscript.

## Data availability statement

All data presented in this manuscript are available for download at our Cellular Lifespan Study shiny app: https://columbia-picard.shinyapps.io/shinyapp-Lifespan_Study, with additional data described in(*28*). The raw RNA sequencing and DNA methylation are available at GEO superseries #<u>GSE179849</u>. Source data and fibroblast images are available at https://doi.org/10.6084/m9.figshare.18441998.v2 and https://doi.org/10.6084/m9.figshare.18444731.v1, respectively. All code r script for data preprocessing is available at https://github.com/gav-sturm/Cellular_Lifespan_Study. Other information is available upon request from the corresponding author.

---

**Figure S1. Evaluation of bioenergetic parameters.** (**A**) Study design: primary human fibroblasts derived from three healthy donors (Donors 1, 2, 3) were cultured under standard conditions or chronically treated with Dexamethasone (Dex, 100 nM) across its lifespan for up to 150 -250 days. Bioenergetics parameters were evaluated every 15-10 days performing the Seahorse Mito Stress Test (Agilent). Green bars highlight the cell volume values the day prior to the test, which were then utilized to normalize the bioenergetic parameters. (**B**) Representative curves of cellular oxygen consumption rate (OCR) and extracellular acidification rate (ECAR) obtained throughout the Seahorse Mito Stress Test (**C**). Bioenergetic parameters that can be derived from ECAR and OCR curves in B. *J*_ATP-Glyc_: ATP production rate derived from glycolysis. *J*_ATP-OxPhos_: ATP production rate derived from OxPhos. O: Oligomycin. FCCP: Carbonyl cyanide-p-trifluoromethoxyphenyl-hydrazon. R/A: Rotenone/Antimycin A.

**Figure S2. Effects of chronic glucocorticoid signaling on basal bioenergetic parameters. (A)** Lifespan trajectories (*left* panel) and lifespan average effects (*right* panel) of Dex treatment on basal *J*_ATP-Glyc_ /20k cells expressed relative to the corresponding control time points for each donor. (**B**) Same as A but for *J*_ATP-OxPhos_/20k cells. (**C**) Same as A but for *J*_ATP-Total_/20k cells. (**D**) Correlation between basal *J*_ATP-Total_ and basal *J*_ATP-Glyc_ (*left* panel) and between basal *J*_ATP-Total and_ basal *J*_ATP-OxPhos_ (*right* panel). n = 3 donors per group, 8-13 timepoints per donor. Lifespan average graphs are mean ± SEM, two-way ANOVA. Correlation graphs show Spearman r and thick lines represent simple linear regression for each group. ** p < 0.01, *** p < 0.001, **** p < 0.0001, ns: not significant. *J*_ATP-Glyc_: ATP production rate derived from glycolysis. *J*_ATP-OxPhos_: ATP production rate derived from OxPhos. *J*_ATP-Total_: algebraic sum of *J*ATP-Glyc and *J*ATP-OxPhos.

**Figure S3. Effects of chronic glucocorticoid signaling on mtDNAcn.** (**A**) Lifespan trajectories of Dex treatment on mtDNA copy number (mtDNAcn). (**B**) Lifespan trajectories (*left* panel) and lifespan average effects (*right* panel) of Dex treatment on mtDNAcn expressed relative to the corresponding control time points for each donor. n = 3 donors per group, 5-8 timepoints per donor. Lifespan average graphs are mean ± SEM, two-way ANOVA. * p < 0.05, ns: not significant.

**Figure S4. Effects of chronic glucocorticoid signaling on gene expression. (A-E)** Heatmaps from Figure 4 shown with complete labelling for gene-specific resolution. (**A**) Heatmaps showing the effect of Dex treatment on the expression of glycolytic genes, expressed as the Log_2_ of fold change (Log_2_FC) of normalized gene expression relative to the corresponding control time point for each donor. (**B**) Same as A but for genes that encodes for the subunits of Complex I, II, III, IV and V (CI, CII, CIII, CIV, and CV) of OxPhos, with mitochondrial-encoded genes marked with ❖. (**C-E**) Same as A, but for genes associated with (**C**) mtDNA replication, (**D**) mtDNA maintenance, and (**E**) mitochondrial biogenesis. n = 3 donors per group, 9-10 timepoints per donor.

**Figure S5. Effects of chronic glucocorticoid signaling on mtDNAcn.** (**A**) Correlation between cell-free mitochondrial DNA (cf-mtDNA) and cell-free nuclear DNA (cf-nDNA). (**B**) Lifespan trajectories (*left* panel) and lifespan average effects (*right* panel) of Dex treatment on cf-mtDNA/cf-nDNA ratio expressed relative to the corresponding control time points for each donor. n = 3 donors per group, 6-10 timepoints per donor. Lifespan average graphs are mean ± SEM, two-way ANOVA. Correlation graphs show Spearman r and thick lines represent simple linear regression for each group. * p < 0.05, ** p < 0.01, **** p < 0.0001, ns: not significant.

**Figure S6. Effects of chronic glucocorticoid signaling on gene expression of age-related cytokines.** (**A**) Heatmaps showing the effect of Dex treatment on the expression of age-related cytokines, expressed as the Log_2_ of fold change (Log_2_FC) normalized gene expression relative to the corresponding control time points for each donor. (**B**) Lifespan trajectories of gene expression in cells treated with Dex relative to the corresponding control time points for each donor. Thin curves in soft red represents individual cytokines; thick curves in red represent the average of all cytokines evaluated; thick lines in gray represent the control level. (**C**) Lifespan trajectories (*left* panel) and average effects (*right* panel) of Dex treatment on *Stc1* gene (most upregulated gene) expression expressed relative to the corresponding control time point for each donor. (**D**) Same as C but for *Lum* gene (most downregulated gene). n = 3 donors per group, 9-10 timepoints per donor. Lifespan average graphs are mean ± SEM, two-way ANOVA. * p < 0.05, **** p < 0.0001, ns: not significant.

**Figure S7. Effects of chronic glucocorticoid signaling on secretion and gene expression of age-related cytokines.** (**A-AB**) Lifespan trajectories (*left* panel) and average effects (*right* panel) of Dex treatment on age-related cytokines levels per mL of culture media (u*pper* panel) and normalized gene expression (*lower* panel), expressed relative to the corresponding control time point for each donor. n = 3 donors per group, 6-10 timepoints per donor. Lifespan average graphs are mean ± SEM, two-way ANOVA. * p < 0.05, ** p < 0.01, *** p < 0.001, **** p < 0.0001, ns: not significant.

**Figure S8. Effects of chronic glucocorticoid signaling on mtDNA instability.** (**A**) Circos plots from mtDNA sequencing and eKLIPse analysis for mtDNA. Red lines represent individual deletion break points that span the sequence contained between its two ends, and line darkness reflects the level of heteroplasmy for each deletion. The blue histograms represent coverage depth along the mtDNA sequence. (**B**) Lifespan trajectories (left panel) and lifespan average effects (right panel) of Dex treatment on the individual deletions count. (**C**) Frequency distribution of deletion lengths for all timepoints across lifespan. (**D**) Frequency of deletions (upper panels) and frequency of deletion break points (lower panels) along the mtDNA sequence. (**E**) Lifespan trajectories (left panel) and lifespan average effects (right panel) of Dex treatment on the deletions that span the D-loop region of the mtDNA. n = 3 donors per group, 2-6 timepoints per donor. Lifespan average graphs are mean ± SEM, two-way ANOVA. ns: not significant; N/A: not applicable.

**Figure S9. Effects of chronic glucocorticoid signaling on cell division.** (**A**) Lifespan trajectories (*left* panel) and average effects (*right* panel) of Dex treatment on doubling rate (divisions/day) expressed relative to the corresponding control time points for each donor. (**B**) Same as A but for *J*_ATP-Total_/cell volume/division. n = 3 donors per group, timepoints per donor: n = 26-36 in A, 8-13 in B. Lifespan average graphs are mean ± SEM, two-way ANOVA. **** p < 0.0001. ns: not significant. *J*_ATP-Total_: algebraic sum of *J*_ATP-Glyc_ and *J*_ATP-OxPhos_. *J*_ATP-Glyc_: ATP production rate derived from glycolysis. *J*_ATP-OxPhos_: ATP production rate derived from OxPhos.

**Figure S10. Effects of chronic glucocorticoid signaling on telomeres.** (**A**) Telomere length across population doublings calculated by the DNAmTL epigenetic clock with linear regressions for each donor of each group (**B**) Telomere length across population doublings calculated by the principal components (PC)DNAmTL epigenetic clock with linear regressions for each donor of each group (*left* panel), and telomere shortening rate inferred by linear mixed model along the whole lifespan (*right* panel). (**C**) Heatmaps showing the effect of Dex treatment on the expression of telomere maintenance-associated genes, expressed as the Log_2_ fold change (Log_2_FC) relative to the corresponding control time point for each donor. (**D**) Average effect of Dex treatment shown in C. Each datapoint represents the gene average of the Log_2_FC values throughout the entire lifespan of the three donors. n = 3 donors per group, timepoints per donor: n = 4-13 in A-B, n = 9-10 in C. Thin lines in correlation graphs show linear regression for each donor, Pearson r correlation. Bar graphs are mean ± SEM, Satterthwaite method in B, and one-sample t-test different than 0 in D. * p < 0.05, ns: not significant.

**Figure S11. Effects of chronic glucocorticoid signaling on aging epigenetic clocks.** (**A**) Epigenetic age across population doublings calculated by the DNAm Hannum epigenetic clock (*left* panel) and the principal components (PC) version of it (*middle* panel), with linear regressions for each donor of each group, and the epigenetic aging rate inferred by linear mixed model along the whole lifespan (*right* panel). (**B-D**) Same as A but for Skin and Blood, Pan Tissue and GrimAge epigenetic clocks, respectively. n = 3 donors per group, 4-13 timepoints per donor. Thin lines in correlation graphs show linear regression for each donor, Pearson r correlation. Bar graphs are mean ± SEM, Satterthwaite method. * p < 0.05, ns: not significant.

**Figure S12. Effects of chronic glucocorticoid signaling and mitochondrial nutrient uptake inhibitors on basal bioenergetic parameters.** (A) Schematic of mitoNUITs mechanism of action, three pharmacological inhibitors of nutrient uptake in the mitochondria. (B) Lifespan average effects of Dex+mitoNUITs treatment on *J*_ATP-OxPhos_/cell volume. (C) Same as (B) but for *J*_ATP-Glyc_/cell volume. N = 3 donors per group, n=7-11 timepoints per donor. Lifespan average graphs are mean ± SEM, two-way ANOVA. * p < 0.05, ** p < 0.01, *** p < 0.001, **** p < 0.0001. mitoNUITs: Mitochondrial nutrient uptake inhibitors. *J*_ATP-Glyc_: ATP production rate derived from glycolysis. *J*_ATP-OxPhos_: ATP production rate derived from OxPhos

