## Supplementary Material for "Cellular Allostatic Load is linked to Increased Energy Expenditure and Accelerated Biological Aging"

**Figure S1**

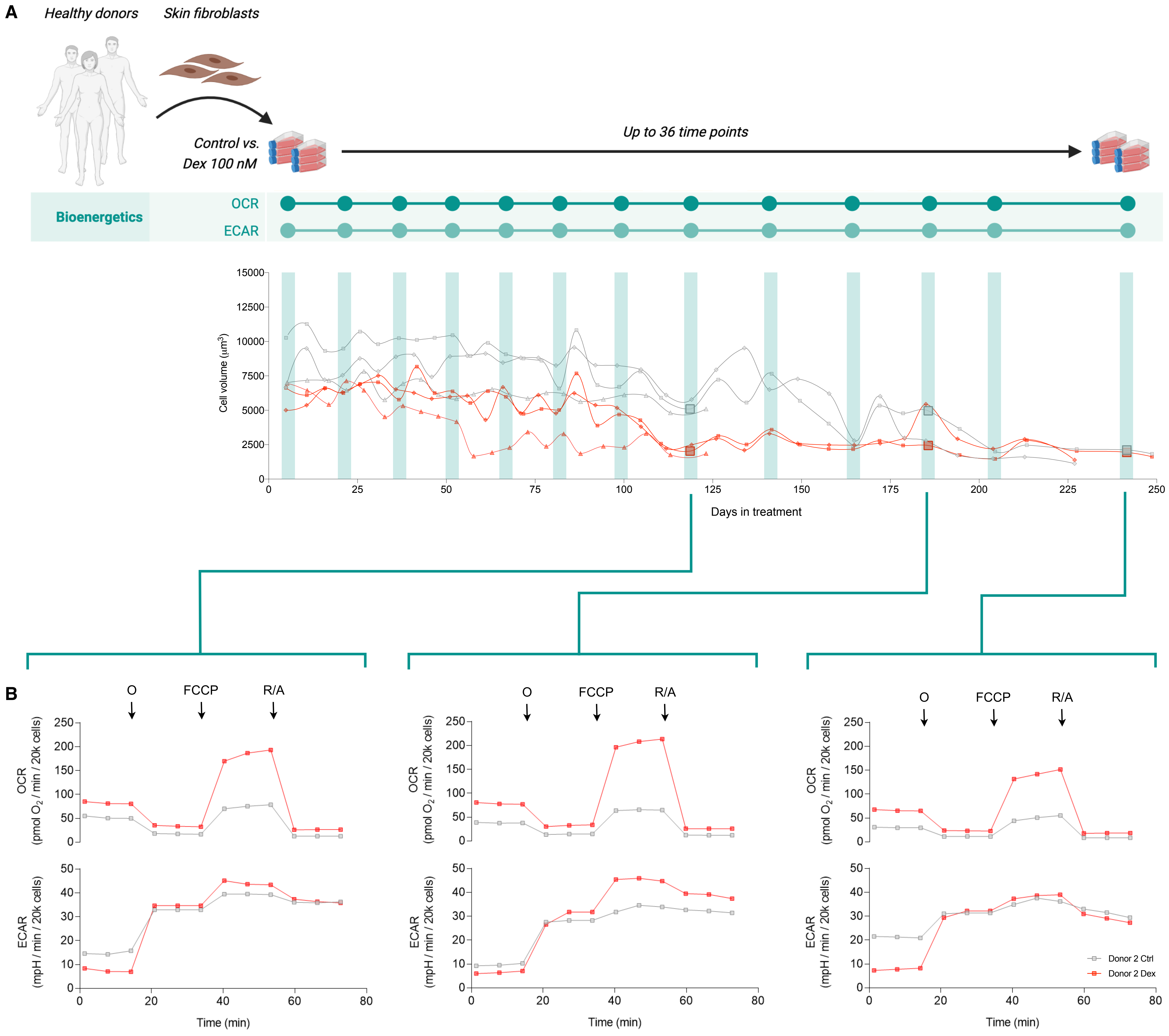


**C**

| | |
|---|---|
| **Basal $J_{ATP\text{-}Glyc}$** | ATP production rate derived from glycolysis under basal conditions |
| **Basal $J_{ATP\text{-}OxPhos}$** | ATP production rate derived from OxPhos under basal conditions |
| **Basal $J_{ATP\text{-}Total}$** | ATP production rate derived from both glycolysis and OxPhos under basal conditions |
| **Compensatory $J_{ATP\text{-}Glyc}$** | ATP production rate derived from glycolysis under inhibition of OxPhos ATP production |
| **Compensatory glycolytic capacity** | Compensatory $J_{ATP\text{-}Glyc}$ expressed as a percentage of basal $J_{ATP\text{-}Total}$ |
| **Maximal $J_{ATP\text{-}OxPhos}$** | ATP production rate derived from OxPhos under maximal respiratory activity |
| **Spare $J_{ATP\text{-}OxPhos}$** | ATP production rate derived from OxPhos under maximal respiratory activity without accounting for the basal $J_{ATP\text{-}OxPhos}$ |
| **Spare OxPhos capacity** | ATP production rate derived from OxPhos expressed as a percentage of basal $J_{ATP\text{-}OxPhos}$ |

**Figure S2**

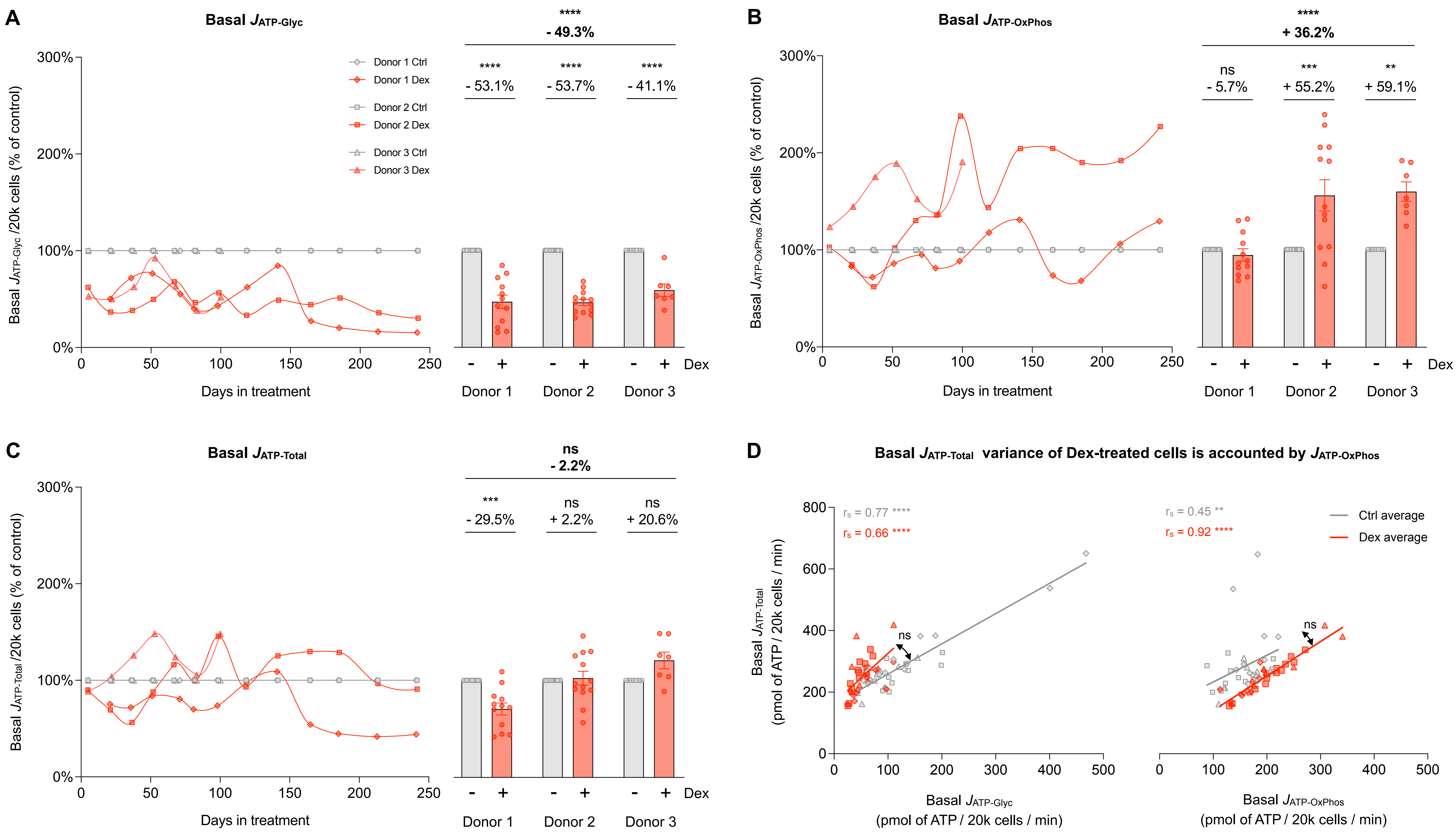

A
Basal $J_{ATP-Glyc}$
Basal $J_{ATP-Glyc}$ /20k cells (% of control)
Donor 1 Ctrl
Donor 1 Dex
Donor 2 Ctrl
Donor 2 Dex
Donor 3 Ctrl
Donor 3 Dex
Days in treatment
****
- 49.3%
- 53.1%
- 53.7%
- 41.1%
Dex
Donor 1
Donor 2
Donor 3
B
Basal $J_{ATP-OxPhos}$
Basal $J_{ATP-OxPhos}$ /20k cells (% of control)
+ 36.2%
ns
- 5.7%
***
+ 55.2%
**
+ 59.1%
C
Basal $J_{ATP-Total}$
Basal $J_{ATP-Total}$ /20k cells (% of control)
- 2.2%
- 29.5%
+ 2.2%
+ 20.6%
D
Basal $J_{ATP-Total}$ variance of Dex-treated cells is accounted by $J_{ATP-OxPhos}$
$r_s$ = 0.77 ****
$r_s$ = 0.66 ****
$r_s$ = 0.45 **
$r_s$ = 0.92 ****
Ctrl average
Dex average
Basal $J_{ATP-Total}$ (pmol of ATP / 20k cells / min)
Basal $J_{ATP-Glyc}$ (pmol of ATP / 20k cells / min)
Basal $J_{ATP-OxPhos}$ (pmol of ATP / 20k cells / min)

**Figure S3**

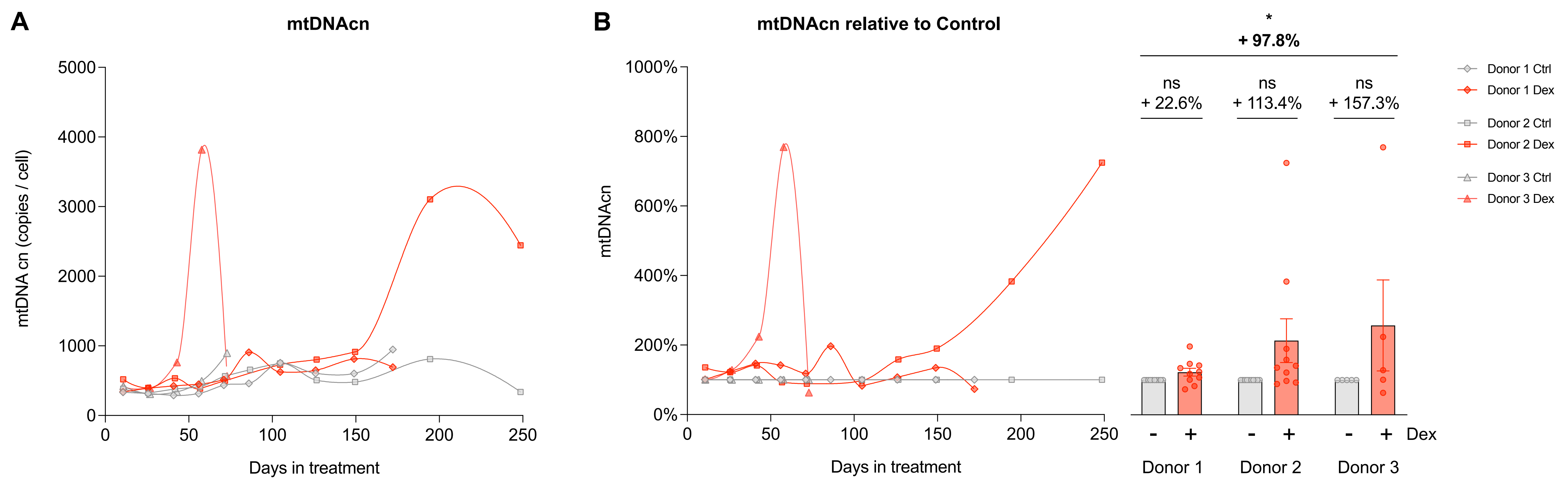
A
mtDNAcn
mtDNA cn (copies / cell)
5000
4000
3000
2000
1000
0
0
50
100
150
200
250
Days in treatment
B
mtDNAcn relative to Control
mtDNAcn
1000%
800%
600%
400%
200%
0%
0
50
100
150
200
250
Days in treatment
*
+ 97.8%
ns
+ 22.6%
ns
+ 113.4%
ns
+ 157.3%
- + - + - + Dex
Donor 1
Donor 2
Donor 3
Donor 1 Ctrl
Donor 1 Dex
Donor 2 Ctrl
Donor 2 Dex
Donor 3 Ctrl
Donor 3 Dex

### Figure S4

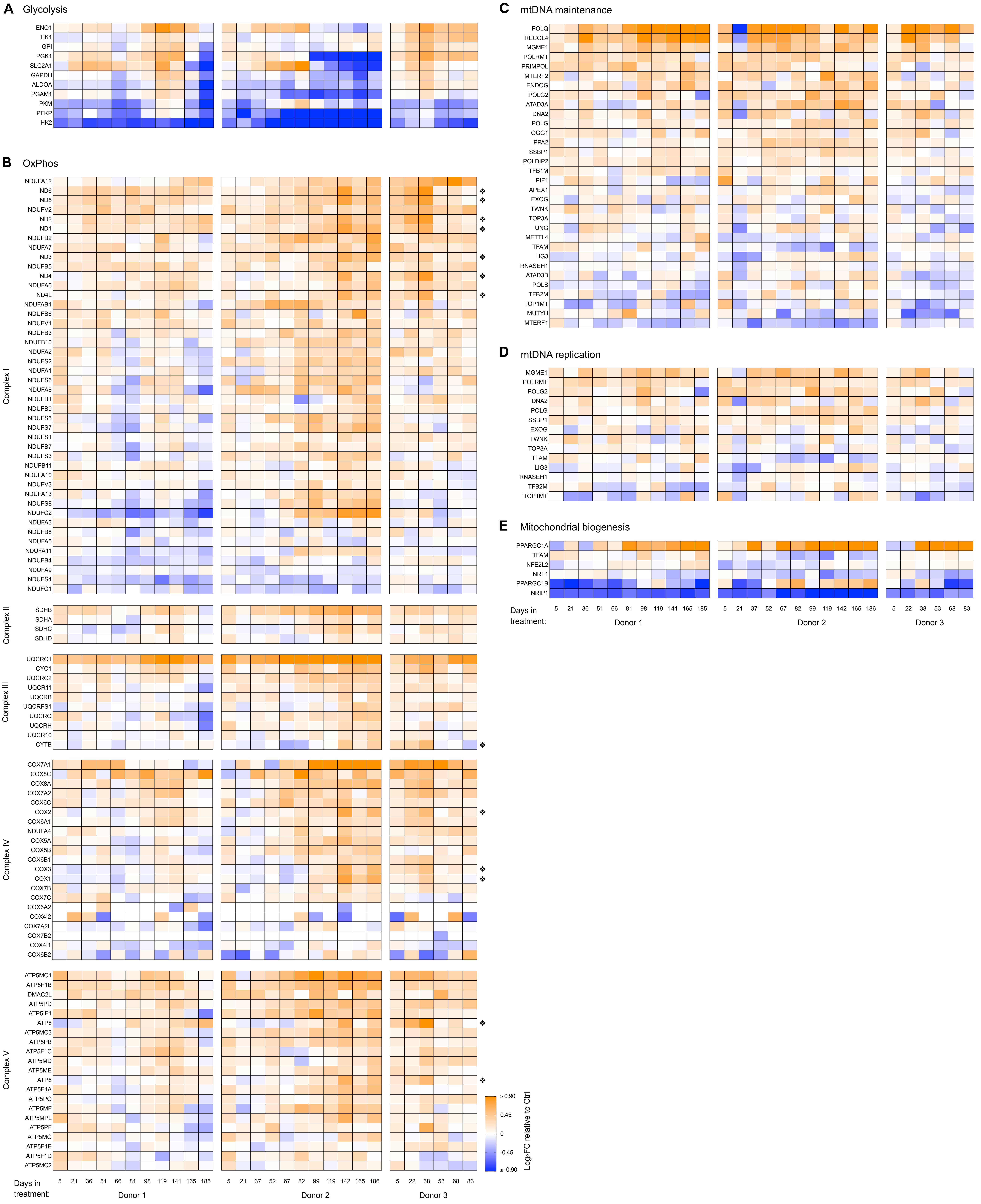
A Glycolysis
ENO1
HK1
GPI
PGK1
SLC2A1
GAPDH
ALDOA
PGAM1
PKM
PFKP
HK2
B OxPhos
Complex I
NDUFA12
ND6
ND5
NDUFV2
ND2
ND1
NDUFB2
NDUFA7
ND3
NDUFB5
ND4
NDUFA6
ND4L
NDUFAB1
NDUFB6
NDUFV1
NDUFB3
NDUFB10
NDUFA2
NDUFS2
NDUFA1
NDUFS6
NDUFA8
NDUFB1
NDUFB9
NDUFS5
NDUFS7
NDUFS1
NDUFB7
NDUFS3
NDUFB11
NDUFA10
NDUFV3
NDUFA13
NDUFS8
NDUFC2
NDUFA3
NDUFB8
NDUFA5
NDUFA11
NDUFB4
NDUFA9
NDUFS4
NDUFC1
Complex II
SDHB
SDHA
SDHC
SDHD
Complex III
UQCRC1
CYC1
UQCRC2
UQCR11
UQCRB
UQCRFS1
UQCRQ
UQCRH
UQCR10
CYTB
Complex IV
COX7A1
COX8C
COX8A
COX7A2
COX6C
COX2
COX6A1
NDUFA4
COX5A
COX5B
COX6B1
COX3
COX1
COX7B
COX7C
COX6A2
COX4I2
COX7A2L
COX7B2
COX4I1
COX6B2
Complex V
ATP5MC1
ATP5F1B
DMAC2L
ATP5PD
ATP5IF1
ATP8
ATP5MC3
ATP5PB
ATP5F1C
ATP5MD
ATP5ME
ATP6
ATP5F1A
ATP5PO
ATP5MF
ATP5MPL
ATP5PF
ATP5MG
ATP5F1E
ATP5F1D
ATP5MC2
Days in treatment:
5 21 36 51 66 81 98 119 141 165 185
Donor 1
5 21 37 52 67 82 99 119 142 165 186
Donor 2
5 22 38 53 68 83
Donor 3
≥ 0.90
0.45
0
-0.45
≤ -0.90
Log2FC relative to Ctrl
C mtDNA maintenance
POLQ
RECQL4
MGME1
POLRMT
PRIMPOL
MTERF2
ENDOG
POLG2
ATAD3A
DNA2
POLG
OGG1
PPA2
SSBP1
POLDIP2
TFB1M
PIF1
APEX1
EXOG
TWNK
TOP3A
UNG
METTL4
TFAM
LIG3
RNASEH1
ATAD3B
POLB
TFB2M
TOP1MT
MUTYH
MTERF1
D mtDNA replication
MGME1
POLRMT
POLG2
DNA2
POLG
SSBP1
EXOG
TWNK
TOP3A
TFAM
LIG3
RNASEH1
TFB2M
TOP1MT
E Mitochondrial biogenesis
PPARGC1A
TFAM
NFE2L2
NRF1
PPARGC1B
NRIP1
Days in treatment:
5 21 36 51 66 81 98 119 141 165 185
Donor 1
5 21 37 52 67 82 99 119 142 165 186
Donor 2
5 22 38 53 68 83
Donor 3

**Figure S5**

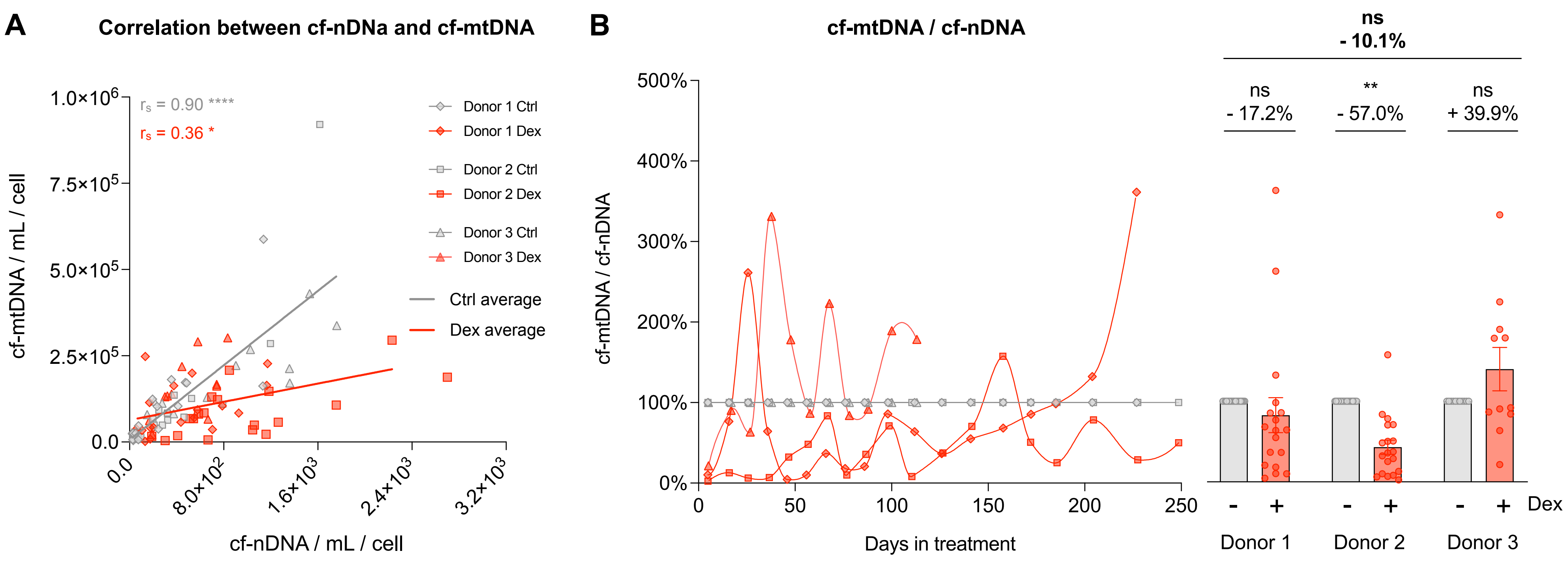
A
Correlation between cf-nDNa and cf-mtDNA
$r_s$ = 0.90 ****
$r_s$ = 0.36 *
Donor 1 Ctrl
Donor 1 Dex
Donor 2 Ctrl
Donor 2 Dex
Donor 3 Ctrl
Donor 3 Dex
Ctrl average
Dex average
cf-mtDNA / mL / cell
cf-nDNA / mL / cell
B
cf-mtDNA / cf-nDNA
ns
- 10.1%
ns
- 17.2%
**
- 57.0%
ns
+ 39.9%
cf-mtDNA / cf-nDNA
Days in treatment
Dex
Donor 1
Donor 2
Donor 3

**Figure S6**

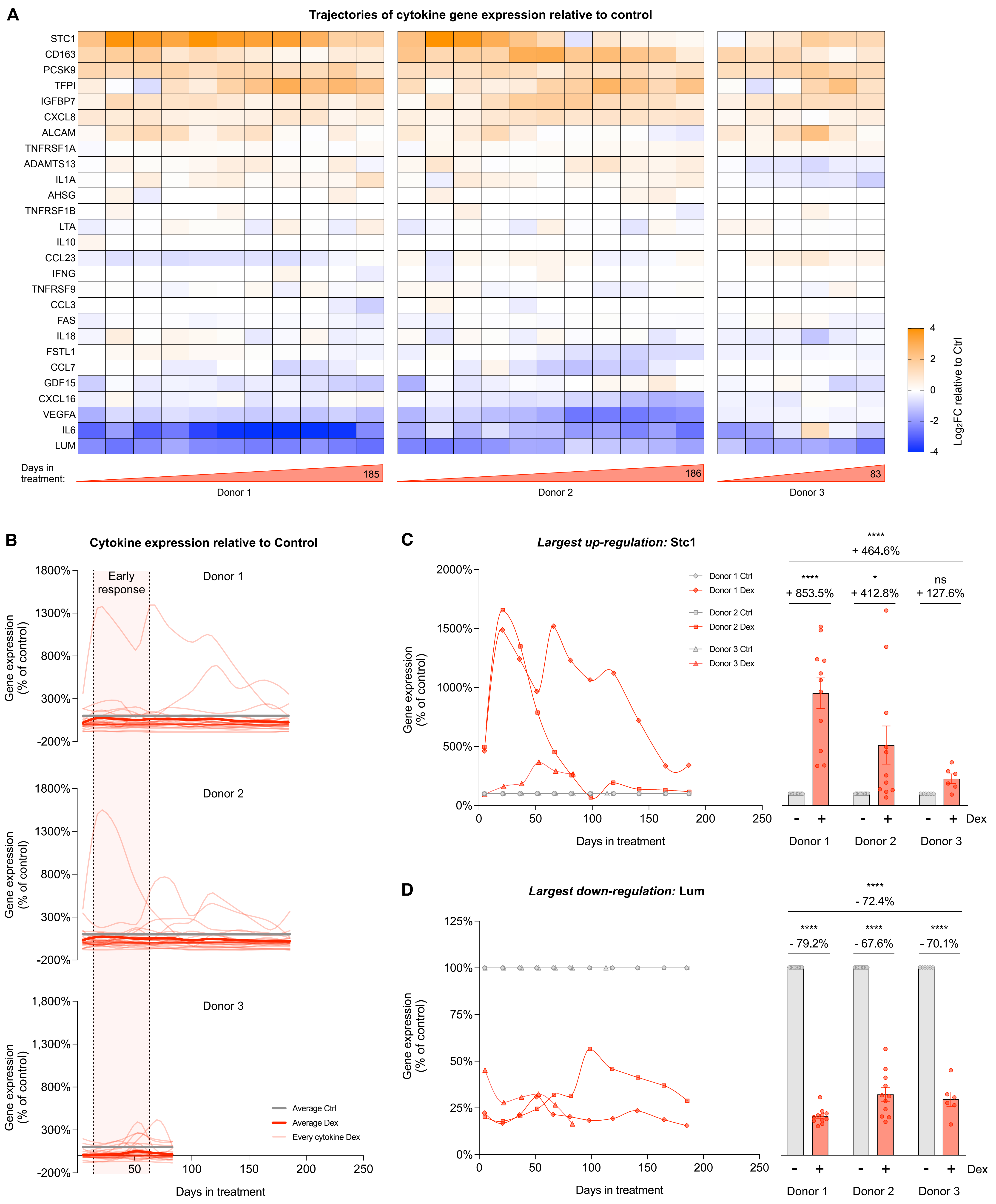
A
Trajectories of cytokine gene expression relative to control
STC1
CD163
PCSK9
TFPI
IGFBP7
CXCL8
ALCAM
TNFRSF1A
ADAMTS13
IL1A
AHSG
TNFRSF1B
LTA
IL10
CCL23
IFNG
TNFRSF9
CCL3
FAS
IL18
FSTL1
CCL7
GDF15
CXCL16
VEGFA
IL6
LUM
Days in treatment:
185
186
83
Donor 1
Donor 2
Donor 3
Log2FC relative to Ctrl
B
Cytokine expression relative to Control
Early response
Donor 1
Donor 2
Donor 3
Gene expression (% of control)
Average Ctrl
Average Dex
Every cytokine Dex
Days in treatment
C
Largest up-regulation: Stc1
Donor 1 Ctrl
Donor 1 Dex
Donor 2 Ctrl
Donor 2 Dex
Donor 3 Ctrl
Donor 3 Dex
+ 464.6%
+ 853.5%
+ 412.8%
ns
+ 127.6%
Dex
Days in treatment
D
Largest down-regulation: Lum
- 72.4%
- 79.2%
- 67.6%
- 70.1%
Days in treatment

**Figure S7**

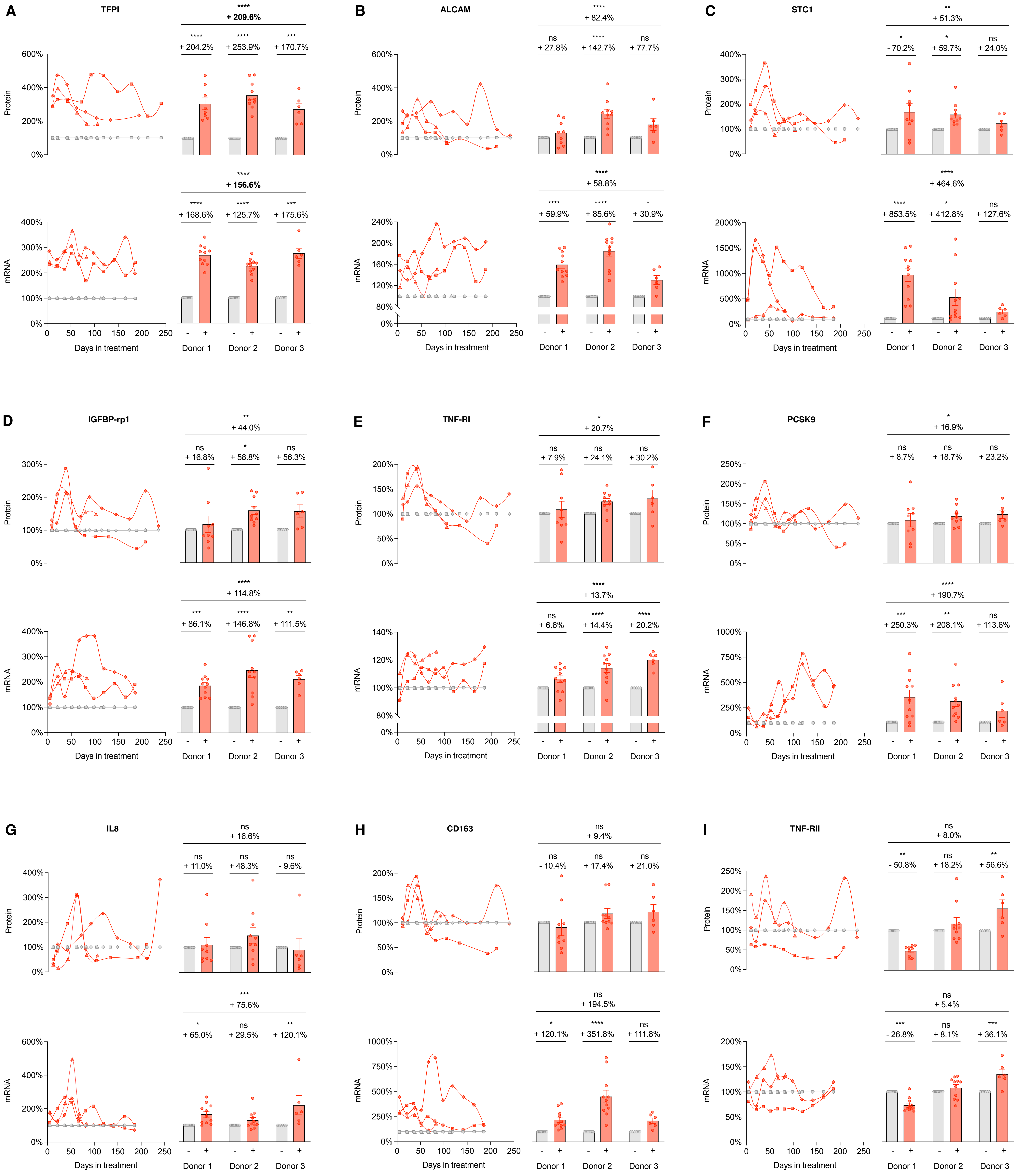

A
TFPI
****
+ 209.6%
****
+ 204.2%
****
+ 253.9%
***
+ 170.7%
Protein
****
+ 156.6%
****
+ 168.6%
****
+ 125.7%
***
+ 175.6%
mRNA
Days in treatment
Donor 1
Donor 2
Donor 3
B
ALCAM
****
+ 82.4%
ns
+ 27.8%
****
+ 142.7%
ns
+ 77.7%
Protein
****
+ 58.8%
****
+ 59.9%
****
+ 85.6%
*
+ 30.9%
mRNA
Days in treatment
Donor 1
Donor 2
Donor 3
C
STC1
**
+ 51.3%
*
- 70.2%
*
+ 59.7%
ns
+ 24.0%
Protein
****
+ 464.6%
****
+ 853.5%
*
+ 412.8%
ns
+ 127.6%
mRNA
Days in treatment
Donor 1
Donor 2
Donor 3
D
IGFBP-rp1
**
+ 44.0%
ns
+ 16.8%
*
+ 58.8%
ns
+ 56.3%
Protein
****
+ 114.8%
***
+ 86.1%
****
+ 146.8%
**
+ 111.5%
mRNA
Days in treatment
Donor 1
Donor 2
Donor 3
E
TNF-RI
*
+ 20.7%
ns
+ 7.9%
ns
+ 24.1%
ns
+ 30.2%
Protein
****
+ 13.7%
ns
+ 6.6%
****
+ 14.4%
****
+ 20.2%
mRNA
Days in treatment
Donor 1
Donor 2
Donor 3
F
PCSK9
*
+ 16.9%
ns
+ 8.7%
ns
+ 18.7%
ns
+ 23.2%
Protein
****
+ 190.7%
***
+ 250.3%
**
+ 208.1%
ns
+ 113.6%
mRNA
Days in treatment
Donor 1
Donor 2
Donor 3
G
IL8
ns
+ 16.6%
ns
+ 11.0%
ns
+ 48.3%
ns
- 9.6%
Protein
***
+ 75.6%
*
+ 65.0%
ns
+ 29.5%
**
+ 120.1%
mRNA
Days in treatment
Donor 1
Donor 2
Donor 3
H
CD163
ns
+ 9.4%
ns
- 10.4%
ns
+ 17.4%
ns
+ 21.0%
Protein
ns
+ 194.5%
*
+ 120.1%
****
+ 351.8%
ns
+ 111.8%
mRNA
Days in treatment
Donor 1
Donor 2
Donor 3
I
TNF-RII
ns
+ 8.0%
**
- 50.8%
ns
+ 18.2%
**
+ 56.6%
Protein
ns
+ 5.4%
***
- 26.8%
ns
+ 8.1%
***
+ 36.1%
mRNA
Days in treatment
Donor 1
Donor 2
Donor 3

**Figure S7 - *Continued (2/3)***

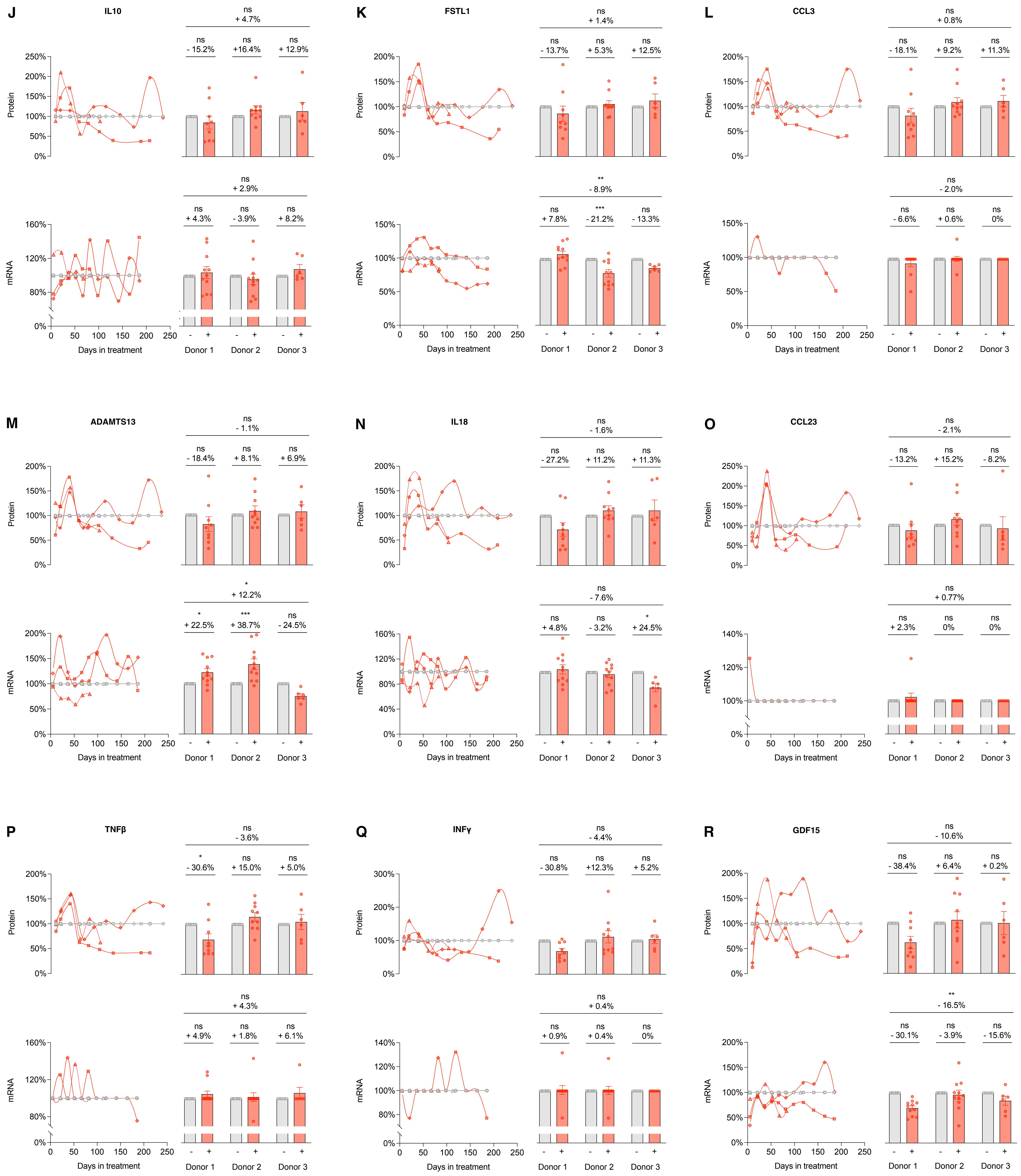

**Figure S7 - *Continued (3/3)***

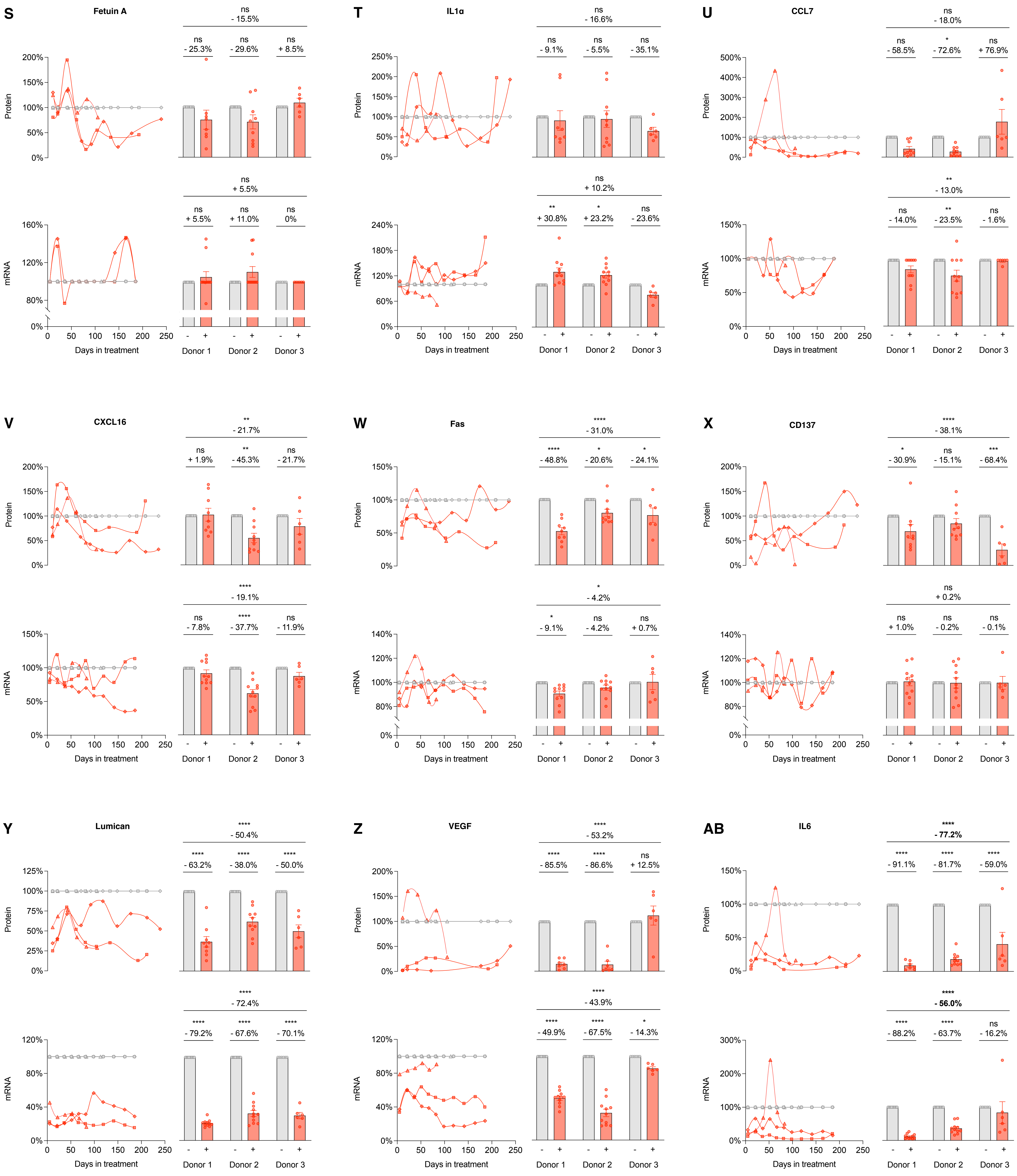

**Figure S8**

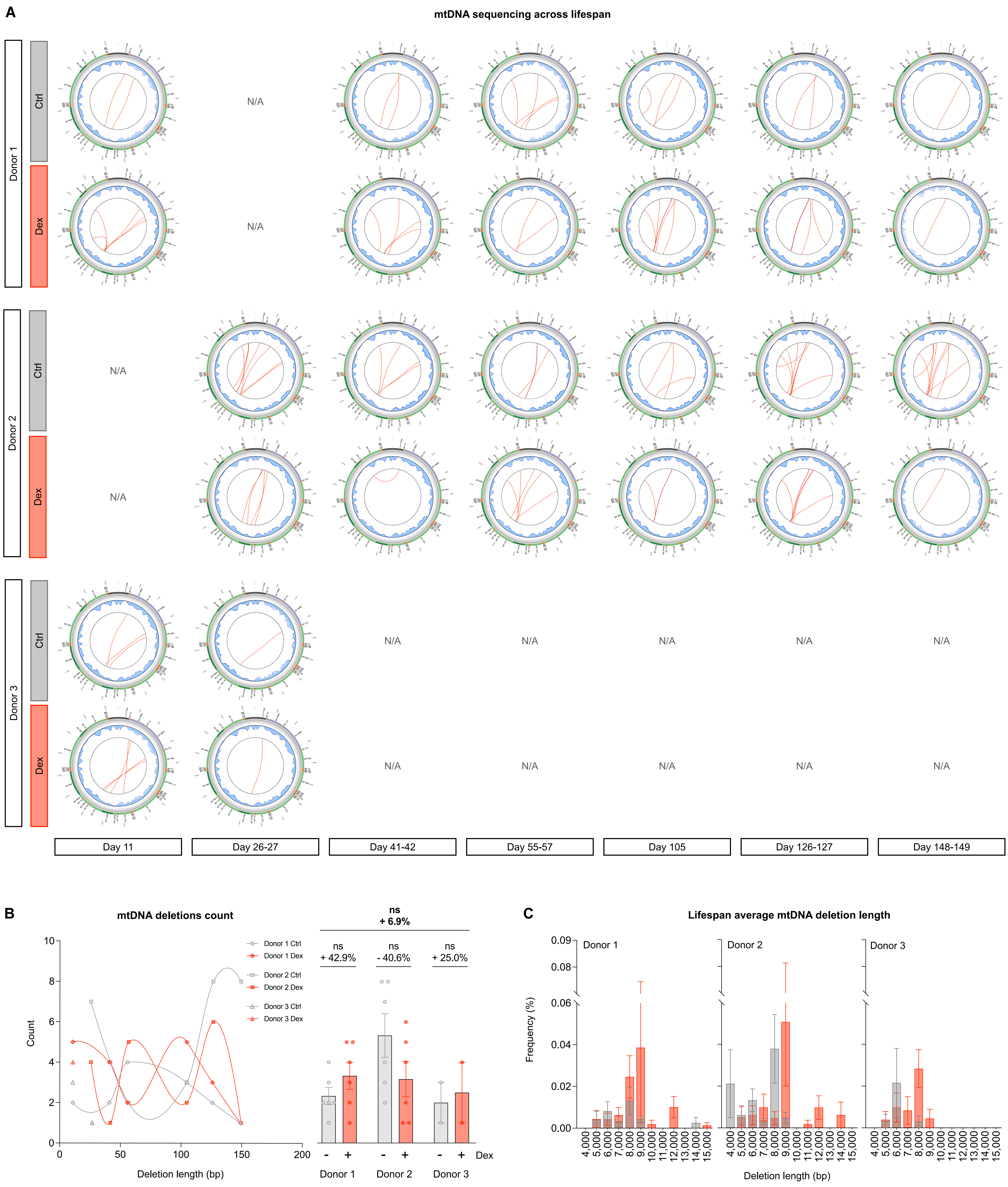
A
mtDNA sequencing across lifespan
Donor 1
Donor 2
Donor 3
Ctrl
Dex
N/A
Day 11
Day 26-27
Day 41-42
Day 55-57
Day 105
Day 126-127
Day 148-149
B
mtDNA deletions count
Count
Deletion length (bp)
0
50
100
150
200
2
4
6
8
10
Donor 1 Ctrl
Donor 1 Dex
Donor 2 Ctrl
Donor 2 Dex
Donor 3 Ctrl
Donor 3 Dex
ns
+ 6.9%
ns
+ 42.9%
ns
- 40.6%
ns
+ 25.0%
-
+
Dex
Donor 1
Donor 2
Donor 3
C
Lifespan average mtDNA deletion length
Frequency (%)
0.00
0.02
0.04
0.06
0.08
0.09
Donor 1
Donor 2
Donor 3
4,000
5,000
6,000
7,000
8,000
9,000
10,000
11,000
12,000
13,000
14,000
15,000
Deletion length (bp)

**Figure S8 - *Continued (2/2)***

D

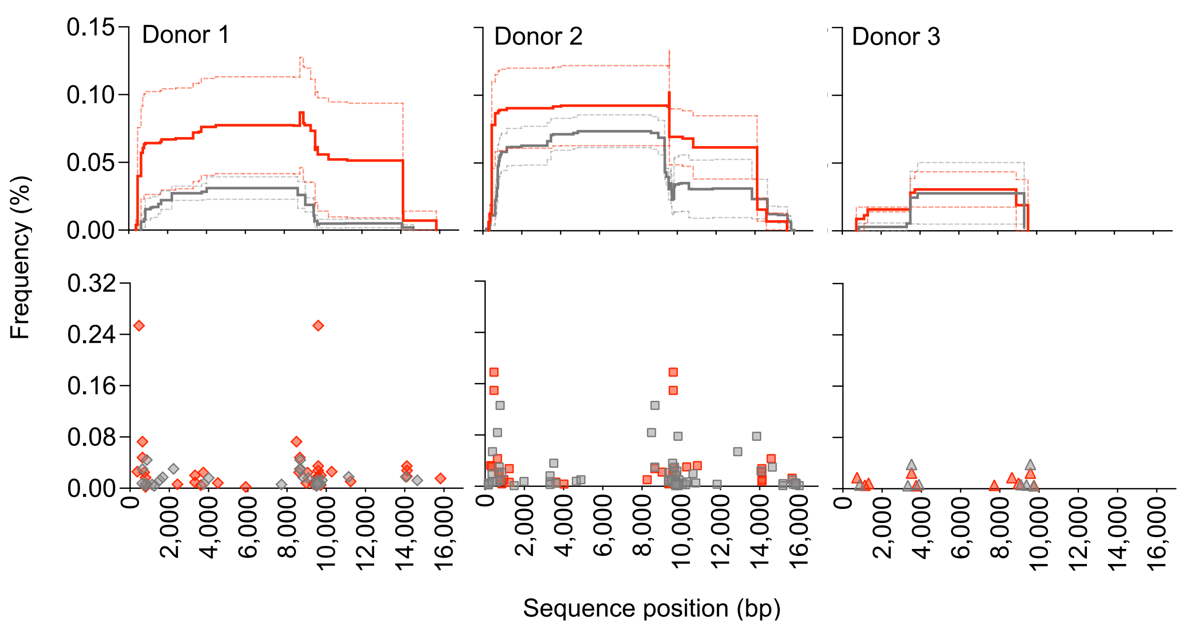


E

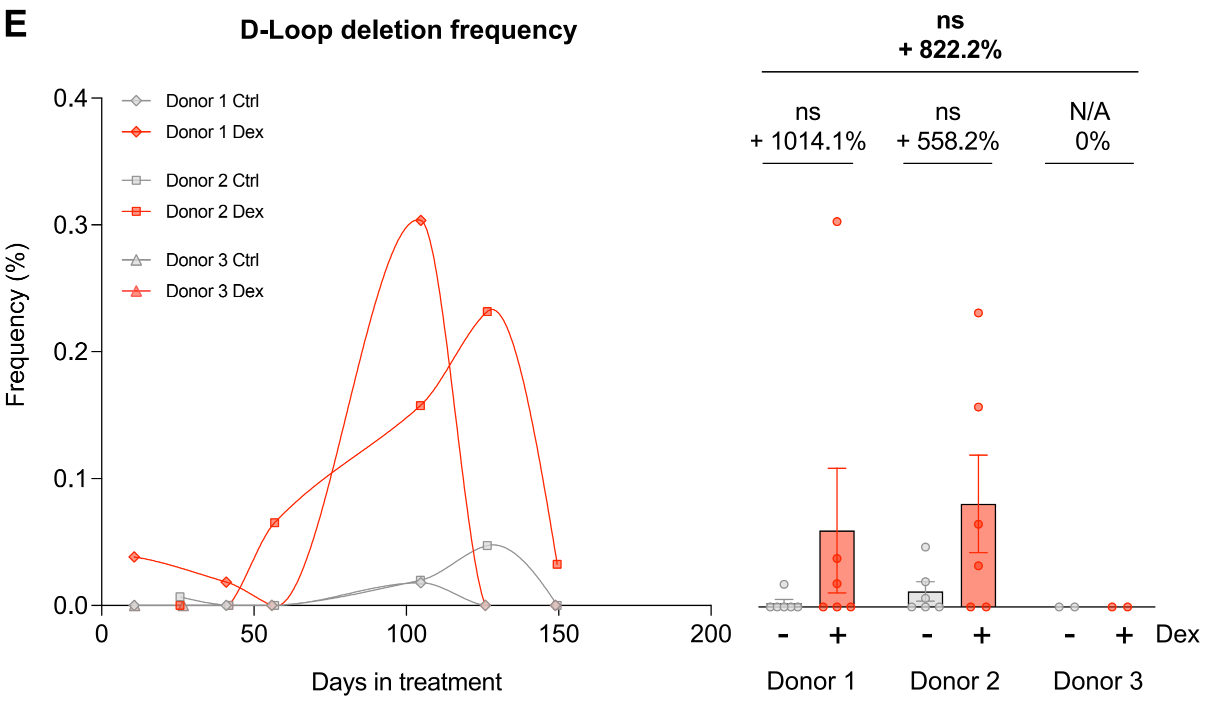

#### Figure S9

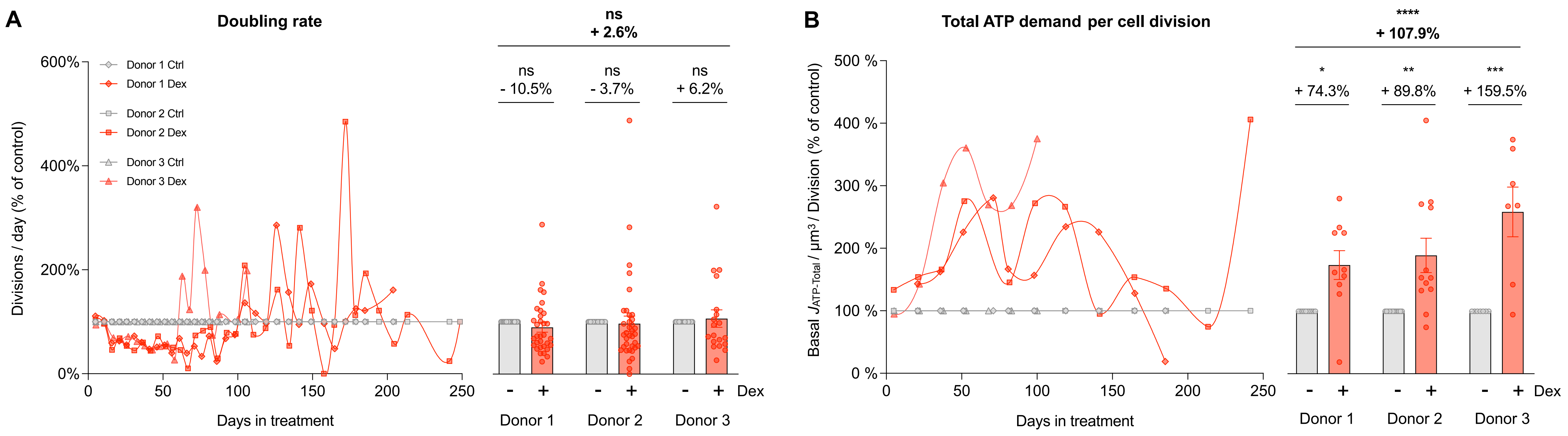
A
Doubling rate
600%
400%
200%
0%
Divisions / day (% of control)
Donor 1 Ctrl
Donor 1 Dex
Donor 2 Ctrl
Donor 2 Dex
Donor 3 Ctrl
Donor 3 Dex
0
50
100
150
200
250
Days in treatment
ns
+ 2.6%
ns
- 10.5%
ns
- 3.7%
ns
+ 6.2%
-
+
Dex
Donor 1
Donor 2
Donor 3
B
Total ATP demand per cell division
500 %
400 %
300 %
200 %
100 %
0 %
Basal $J_{ATP-Total}$ / µm³ / Division (% of control)
Days in treatment
****
+ 107.9%
*
+ 74.3%
**
+ 89.8%
***
+ 159.5%

### Figure S10

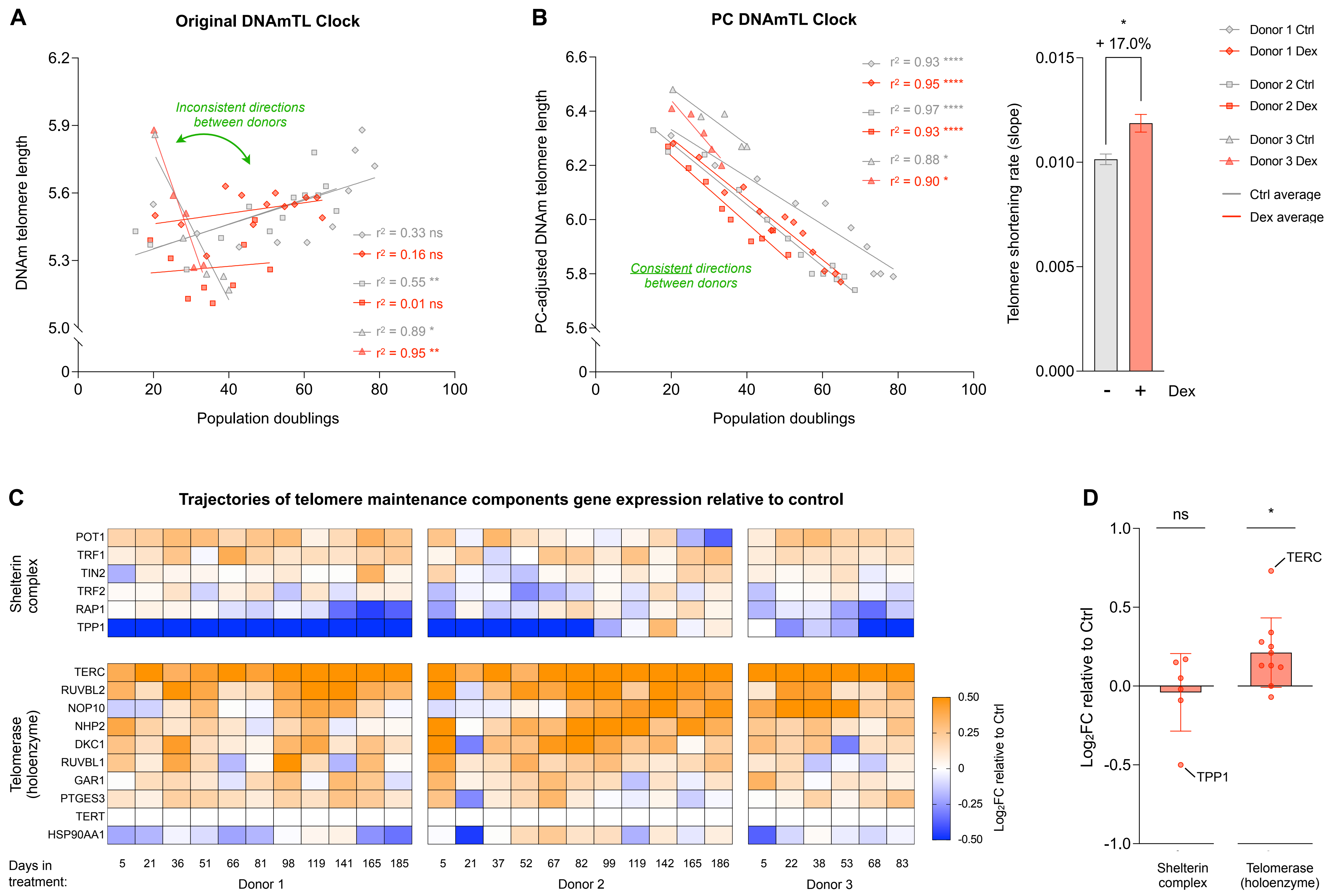

A
Original DNAmTL Clock
DNAm telomere length
Population doublings
Inconsistent directions between donors
r² = 0.33 ns
r² = 0.16 ns
r² = 0.55 **
r² = 0.01 ns
r² = 0.89 *
r² = 0.95 **
B
PC DNAmTL Clock
PC-adjusted DNAm telomere length
Population doublings
Consistent directions between donors
r² = 0.93 ****
r² = 0.95 ****
r² = 0.97 ****
r² = 0.93 ****
r² = 0.88 *
r² = 0.90 *
Telomere shortening rate (slope)
+ 17.0%
Dex
Donor 1 Ctrl
Donor 1 Dex
Donor 2 Ctrl
Donor 2 Dex
Donor 3 Ctrl
Donor 3 Dex
Ctrl average
Dex average
C
Trajectories of telomere maintenance components gene expression relative to control
Shelterin complex
POT1
TRF1
TIN2
TRF2
RAP1
TPP1
Telomerase (holoenzyme)
TERC
RUVBL2
NOP10
NHP2
DKC1
RUVBL1
GAR1
PTGES3
TERT
HSP90AA1
Log2FC relative to Ctrl
Days in treatment:
5 21 36 51 66 81 98 119 141 165 185
Donor 1
5 21 37 52 67 82 99 119 142 165 186
Donor 2
5 22 38 53 68 83
Donor 3
D
Log2FC relative to Ctrl
ns
TERC
TPP1
Shelterin complex
Telomerase (holoenzyme)

**Figure S11**

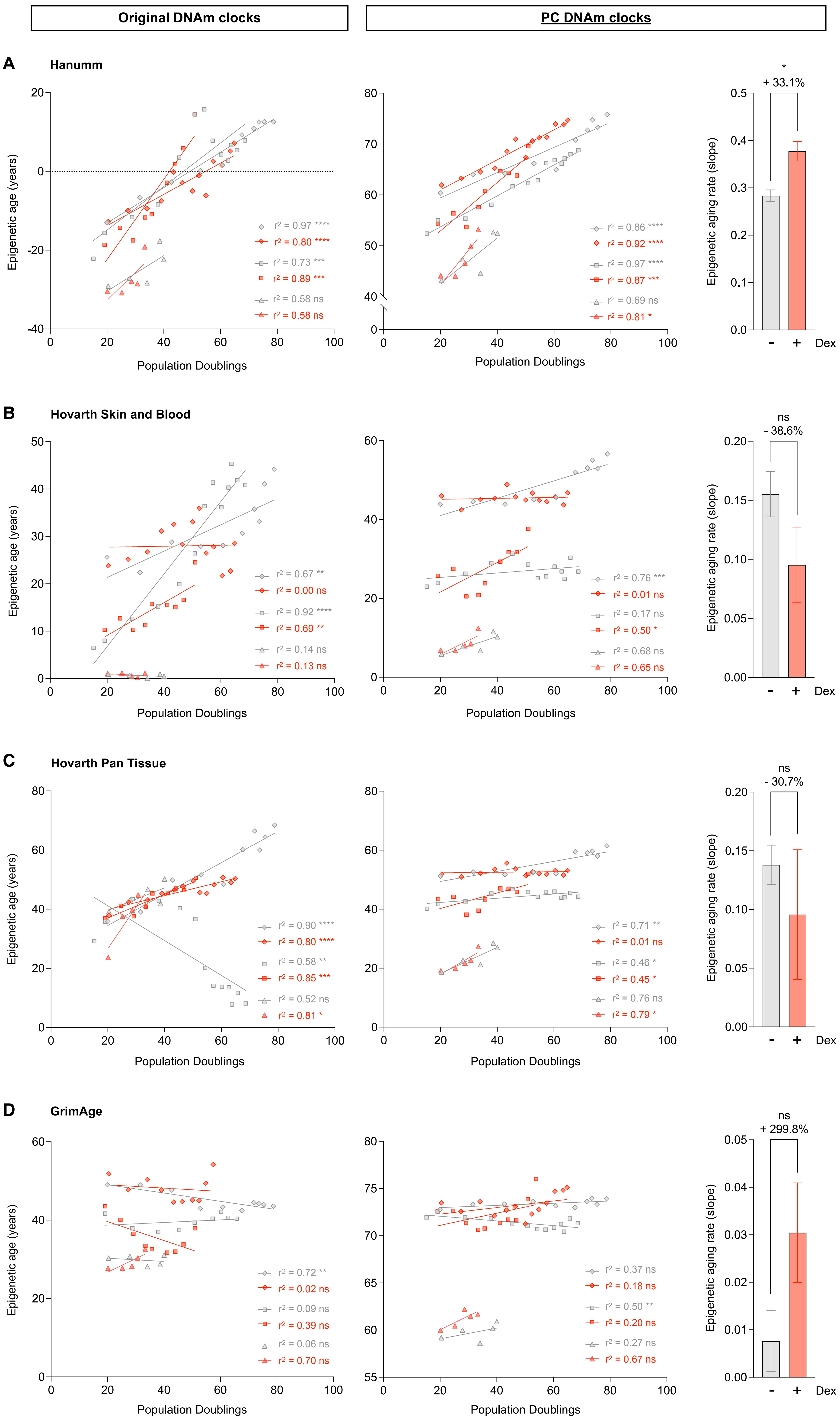

Original DNAm clocks
PC DNAm clocks
A
Hanumm
Epigenetic age (years)
Population Doublings
r² = 0.97 ****
r² = 0.80 ****
r² = 0.73 ***
r² = 0.89 ***
r² = 0.58 ns
r² = 0.58 ns
r² = 0.86 ****
r² = 0.92 ****
r² = 0.97 ****
r² = 0.87 ***
r² = 0.69 ns
r² = 0.81 *
*
+ 33.1%
Epigenetic aging rate (slope)
Dex
B
Hovarth Skin and Blood
r² = 0.67 **
r² = 0.00 ns
r² = 0.92 ****
r² = 0.69 **
r² = 0.14 ns
r² = 0.13 ns
r² = 0.76 ***
r² = 0.01 ns
r² = 0.17 ns
r² = 0.50 *
r² = 0.68 ns
r² = 0.65 ns
ns
- 38.6%
C
Hovarth Pan Tissue
r² = 0.90 ****
r² = 0.80 ****
r² = 0.58 **
r² = 0.85 ***
r² = 0.52 ns
r² = 0.81 *
r² = 0.71 **
r² = 0.01 ns
r² = 0.46 *
r² = 0.45 *
r² = 0.76 ns
r² = 0.79 *
ns
- 30.7%
D
GrimAge
r² = 0.72 **
r² = 0.02 ns
r² = 0.09 ns
r² = 0.39 ns
r² = 0.06 ns
r² = 0.70 ns
r² = 0.37 ns
r² = 0.18 ns
r² = 0.50 **
r² = 0.20 ns
r² = 0.27 ns
r² = 0.67 ns
ns
+ 299.8%

**Figure S12**

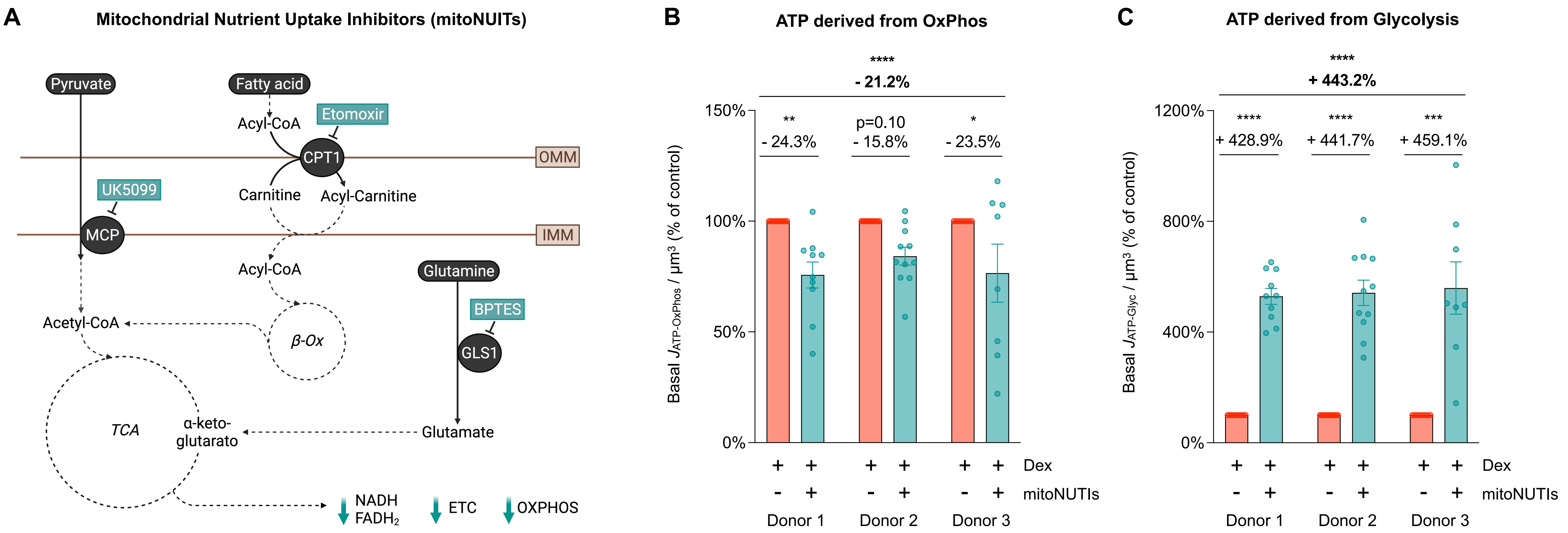
A
Mitochondrial Nutrient Uptake Inhibitors (mitoNUITs)
Pyruvate
Fatty acid
Acyl-CoA
Etomoxir
CPT1
OMM
UK5099
Carnitine
Acyl-Carnitine
MCP
IMM
Acyl-CoA
Glutamine
BPTES
Acetyl-CoA
β-Ox
GLS1
TCA
α-keto-glutarato
Glutamate
NADH
FADH2
ETC
OXPHOS
B
ATP derived from OxPhos
****
- 21.2%
**
- 24.3%
p=0.10
- 15.8%
*
- 23.5%
150%
100%
50%
0%
Basal JATP-OxPhos / µm3 (% of control)
+ + + + + + Dex
- + - + - + mitoNUTIs
Donor 1 Donor 2 Donor 3
C
ATP derived from Glycolysis
****
+ 443.2%
****
+ 428.9%
****
+ 441.7%
***
+ 459.1%
1200%
800%
400%
0%
Basal JATP-Glyc / µm3 (% of control)
+ + + + + + Dex
- + - + - + mitoNUTIs
Donor 1 Donor 2 Donor 3
